# A chimeric, half-life extended lysin with a unique mode of action

**DOI:** 10.64898/2026.05.13.724763

**Authors:** Zehra Visram, Donaat Kestemont, Benedikt Bauer, Renping Qiao, Svetlana Durica-Mitic, Andrea Majoros-Hashempour, Julia Schmidt, Rocío Berdaguer, Karsten Krey, Michele Mutti, Manuel Zerbs, Marina von Freyberg, Lorenzo Corsini, Adriana Badarau

**Affiliations:** BioNTech R&D Austria (Vienna, Austria)

**Keywords:** endolysin, staphylococci, targeted release, enzyme engineering, antibacterial agents, half-life extension

## Abstract

Anti-staphylococcal lytic agents, such as lysostaphin (LSN), a glycyl-glycyl (Gly-Gly) peptidase, have long been considered for the management of chronic and complicated bacterial infections typically resistant to conventional antibiotics, but their use is restricted by poor pharmacokinetic properties. We generated half-life extended lysostaphin constructs by fusing the lysin - either alone or chimerized with an additional enzymatic cysteine, histidine-dependent amidohydrolases/peptidase (CHAP) domain - to the human IgG1 Fc fragment. The Fc-CHAP-LSN constructs retain high potency against *Staphylococcus aureus* and coagulase negative staphylococcal strains *in vitro* and are efficacious in *S. aureus ex vivo* biofilm models and *in vivo* sepsis models. A detailed investigation of the Fc-CHAP-LSN mode of action revealed that upon binding to the bacterial cell, but not in solution, LSN mediates its own release by cleaving the Gly-Gly CHAP-LSN linker. The bactericidal activity of Fc-CHAP-LSN is driven by the LSN-catalyzed and target cell-dependent release of free LSN. The half-life extended lysin acts as a pro-drug, unveiling a novel mechanism of targeted release and an alternative approach to half-life extension.

## Introduction

*Staphylococcus aureus* is a major human pathogen causing skin infections, osteomyelitis, endocarditis, and biofilm-associated device infections (Trobos et al., 2022; Bastien et al., 2023; Kaushik et al., 2024; Tong et al., 2025). The rise of multidrug-resistant strains (e.g. MRSA) has intensified the search for alternative therapeutics. Endolysins and chimeric lysins are attractive candidates due to their rapid bactericidal action, low resistance profile and efficacy in serum or biofilm environments (Schuch et al., 2014; Fischetti, 2018; Guo et al., 2022; Liu et al., 2024). The clinical development of exebacase (CF-301) demonstrated the translational promise of lysins, even though a phase III trial was halted early due to futility (Fowler et al., 2024).

Most endolysins directed against Gram-positive bacteria display a modular architecture, typically composed of one or more enzymatically active domains (EADs) linked to a cell wall-binding domain (CBD) via flexible interdomain linkers (Cui et al., 2025). This modularity enables domain swapping and structural recombination—features that have been increasingly exploited to engineer improved variants with optimized stability, broadened activity, or refined specificity (Gerstmans et al., 2018; Aitken et al., 2025; Hu et al., 2025).

Lysostaphin (LSN), produced by *Staphylococcus simulans*, is a well-characterized glycyl-glycyl endopeptidase that specifically cleaves cross-bridges of the *S. aureus* peptidoglycan. Its narrow but potent activity has made it one of the most intensively studied anti-staphylococcal agents. LSN consists of an N-terminal catalytic Peptidase M23 (PepM23) domain and a C-terminal SH3b CBD, the latter conferring high affinity and species-selective binding to *S. aureus*. This architecture not only facilitates robust lysis of planktonic bacteria but also contributes to strong activity against biofilms, a major challenge in *S. aureus* infections (Schindler and Schuhardt, 1964; Resch et al., 2011; Huang et al., 2013; Grishin et al., 2019b; Shen et al., 2021; CLSI, 2026). Because of its potency, modularity, and established safety profile in preclinical models, LSN frequently serves as a parental scaffold or “building block” for engineering next-generation chimeric lysins (Schindler and Schuhardt, 1964; Resch et al., 2011). Its well-defined domain boundaries also simplify recombination with heterologous domains, such as cysteine, histidine-dependent amidohydrolases/peptidase (CHAP), PepM23 or amidase domains, enabling hybrid constructs with broadened catalytic activity or increased resilience in complex biological matrices (Schmelcher et al., 2011; Li et al., 2021).

However, LSN, and even chimeric lysins comprising LSN domains, show limited *in vivo* availability and fast serum clearance due to their relatively small size (30 – 40 kDa) below renal filtration limit (Maack et al., 1979) and high positive charge (Miner, 2008). LSN half-life extension has been reported in several studies, e.g. by fusion with albumin binding domains (Seijsing et al., 2018; Grishin et al., 2019b) or dimerization (Grishin et al., 2019a), but while serum half-life extension was successful, this came at the expense of lysin potency due to larger size (which decreases the rate of peptidoglycan penetration) and avidity (Sobieraj Anna et al., 2020).

Here we report the half-life extension of LSN and LSN-containing chimeric lysins by fusion to a human immunoglobulin G1 (IgG1) Fc. The Fc-tagged LSN constructs show good activity against *S. aureus* but are inactive against coagulase negative staphylococci. Introducing a CHAP domain to yield Fc-CHAP-LSN leads to retained potency against *S. aureus* and restored activity against coagulase negative strains *in vitro* as well as to high efficacy in *S. aureus ex vivo* biofilm models and *in vivo* sepsis models. A detailed mode of action analysis of Fc-CHAP-LSN revealed that LSN releases itself from the parental molecule at the *S. aureus* cell surface, which drives bactericidal activity of the construct.

## Methods and Materials

### Construct design

Wild-type lysostaphin (LSN_wt_ - Uniprot ID P10547; amino acids 248 to 493) was used for *Escherichia coli* expression. For secretion in Expi293 cells, the sequence was modified by removal of both (S126P and N232Q; LSN_HEK_) or one (S126P; LSN_HEKa_) N-glycosylation sites. The CHAP-LSN chimeras were generated by N-terminally fusing the CHAP domain of a previously described LysM-CHAP (L2-1), (Badarau et al., 2026) by a GG-linker to LSN_HEK_. To enhance secretion, the Y46D mutation was introduced into the CHAP domain (CHAP_opt_). An additional point mutation (K143A) was introduced which improved LSN release, resulting in variant CHAP_opt_var_. Constructs used for display experiments were fused to a GSG linker, a C-terminal Myc tag, and the platelet-derived growth factor receptor (PDGFR) transmembrane domain (UniProt ID P09619, residues 512–562). Catalytic inactivation of the CHAP domain (CHAP*) was achieved by substituting the cysteine at position 27 with serine. Inactivation of the LSN was achieved by the substitution of histidine at position 82 to alanine (H82A) or histidine at position 113 with alanine (H113A) substitutions resulting in LSN*.

*E.* coli codon-optimized LSN constructs (*^EC^*LSN_wt_ and *^EC^*LSN_HEK_) were cloned into pET-29b (Novagen) under control of the IPTG (isopropyl β-D-1-thiogalactopyranoside)-inducible T7 promoter and expressed in *E. coli*. Lysin sequences were codon-optimized for *E. coli* expression and equipped with an N-terminal hexahistidine tag followed by a Human Rhinovirus 3C (HRV 3C) protease cleavage site (SAGLEVLFQGP).

For secretion from Expi293 cells, lysin sequences were codon-optimized for human expression and cloned into pcDNA3.4 (Thermo Fisher Scientific) under the constitutive CMV promoter. Constructs contained a 5′ Kozak sequence followed by a murine Ig κ light-chain signal peptide (mutant A2: MDMRAPAGIFGFLLVLFPGYRS) (Kober et al., 2013), a TG dipeptide to enhance signal peptide cleavage, an N-terminal hexahistidine tag, and a HRV 3C protease cleavage site. CHAP-LSN constructs were fused to the C-terminus of the human IgG1 Fc domain (amino acids 218-449 relative to UniProt-ID: P0DOX5) containing the C222S mutation. The Fc and CHAP domains were connected via a helical linker with eight repeats (AEAAAKEAAAKEAAAKEAAAKALEAEAAAKEAAAKEAAAKEAAAKA - L1) or a helical linker with 14 repeats followed by a 12-amino-acid serum-stable peptide derived from previously reported linkers (AEAAAKEAAAKEAAAKEAAAKEAAAKEAAAKEAAAKALEAEAAAKEAAAKEAAAKEAAAKEAAAKEAAAKEAAAKAPKETVSLNEESN, L2; or AEAAAKEAAAKEAAAKEAAAKEAAAKEAAAKEAAAKALEAEAAAKEAAAKEAAAKEAAAKEAAAKEAAAKEAAAKAKPETVSGNEKSN, L2a) (Badarau et al., 2026). The alanine scanning mutants were derived from Fc-L2a-CHAP_opt_-LSN_HEK_, constructs were generated in which individual glycine residues were substituted within either the CHAP domain (CHAP_opt-G85A,_ CHAP_opt-G86A,_ CHAP_opt-G87A,_ CHAP_opt-G115A,_ CHAP_opt-G116A_ or CHAP_opt-G117A_) or LSN (LSN_HEK-G62A_, LSN_HEK-G63A_ or LSN_HEK-G64A_). If LSN_HEKa_ was fused to a Fc-domain (N- or C-terminally), L1 was used as a linker.

### Protein production and quality control

#### Protein expression and immobilized metal ion affinity Chromatography (IMAC) purification

Fc-L1-CHAP-LSN_HEK_, Fc-L1-CHAP*-LSN_HEK_, Fc-L1-CHAP_opt_-LSN_HEKa_, Fc-L1-CHAP-LSN*_HEK_, Fc-L1-CHAP_opt_-LSN_HEK_, Fc-L2a-CHAP_opt_var_-LSN_HEK_, Fc-L2-CHAP_opt_-LSN_HEK_ and, Fc-L2a-CHAP_opt_-LSN_HEK_ were purified from Expi293 supernatants. Cultures (80 - 800 mL) of Expi293 cells (Thermo Fisher Scientific) were grown with shaking (120 rpm) at 37 °C and 5% CO_2_ in Expi293 medium according to the manufacturer’s instructions. Transfections were performed using the Expifectamine transfection kit (Thermo Fisher Scientific, A14524) and Optiplex (Thermo Fisher Scientific, A4096801) according to the manufacturer’s instructions and at a final DNA concentration of 1 µg/mL. Enhancer was added after 24 hours (h) and supernatants were harvested 72 h post transfection by centrifugation (500 × g, 30 min). Supernatants were clarified by a second high-speed centrifugation (15000 × g, 30 min, 4°C). Cleared supernatants were adjusted to IMAC loading conditions by supplementation with NaCl (300 mM final), Imidazole (20 mM final) and dithiothreitol (DTT, 1 mM final). Benzonase (Merck Millipore, 70746-3, 0.01% [v/v] final) was added to minimize nucleic acid contamination prior to chromatography. Proteins were purified directly from clarified culture supernatants using Ni²⁺-charged HisTrap HP columns (5 mL; Cytiva) on an ÄKTA pure chromatography system. Columns were equilibrated in binding buffer (50 mM MES pH 6.5, 300 mM NaCl, 1 mM CaCl_2_, 5% Glycerol, 50 mM Arginine, 20 mM Imidazole [pH 6.5], 2 mM DTT). Samples were loaded at 1.5 mL/min, followed by washing the column with binding buffer until baseline absorbance was reached. Bound proteins were eluted using step gradient changes to 20% and 80% of elution buffer (50 mM MES pH 6.5, 300 mM NaCl, 1 mM CaCl_2_, 5% Glycerol, 50 mM Arginine, 1 M Imidazole [pH 6.5], 2 mM DTT). Elution fractions were analyzed by sodium dodecyl sulfate–polyacrylamide gel electrophoresis (SDS-PAGE) followed by Coomassie staining and fractions containing the target protein were pooled. LSN_HEK_, LSN_HEKa_, Fc-L1-LSN_HEKa_ and LSN_HEKa_-L1-Fc were purified via IMAC but with 20 mM HEPES, 300 mM NaCl, 20 mM Imidazole (pH 7), 1mM DTT as binding buffer and 20 mM HEPES, 300 mM NaCl, 1 M Imidazole (pH 7), 1 mM DTT as elution buffer. Elution fractions were analyzed by SDS–PAGE followed by Coomassie staining and fractions containing the target protein were pooled. Pooled eluates were supplemented with 1 mM DTT and HRV 3C protease (produced in-house) at a 1:100 protease:substrate molar ratio, followed by dialysis against 300 volumes of formulation buffer (20 mM HEPES [pH 7.0], 150 mM NaCl). Samples were sterile filtered (0.2 µm cutoff filters), aliquoted and frozen at -80 °C for long-term storage. Protein concentrations were determined by UV absorbance using molar extinction coefficients calculated from the amino acid sequence of each lysin variant using the Geneious software.

#### Size exclusion chromatography (SEC)

Pooled eluates of proteins (Fc-L1-CHAP-LSN_HEK_, Fc-L1-CHAP*-LSN_HEK_, Fc-L1-CHAP_opt_-LSN_HEKa_, Fc-L1-CHAP-LSN*_HEK_, Fc-L1-CHAP_opt_-LSN_HEK_, LSN_HEK_, Fc-L2a-CHAP_opt_var_- LSN_HEK_, Fc-L2-CHAP_opt_-LSN_HEK_) from IMAC purification were concentrated for size-exclusion chromatography on an ÄKTA pure 25 system (Cytiva) using Superdex 200 pg, 16/600 HiLoad column. The column was equilibrated in 50 mM MES (pH 6.5), 300 mM NaCl, 1 mM CaCl_2_, 5% Glycerol, 50 mM Arginine, 2 mM DTT. Protein samples were clarified prior to injection by centrifugation and eluted isocratically at 0.5 mL/min. Elution fractions were analyzed by SDS–PAGE followed by Coomassie staining and monomer fractions containing the target protein were pooled and supplemented with HRV 3C protease (produced in-house) at a 1:100 protease:substrate molar ratio, followed by overnight incubation at 4 °C. Samples were sterile-filtered (0.2 µm cutoff filters), aliquoted and frozen at -80 °C for long-term storage. Protein concentrations were determined by UV absorbance using molar extinction coefficients calculated from the amino acid sequence of each lysin variant using the Geneious software (Dotmatics).

#### Protein purification for lysostaphin (^EC^LSN_WT_ and ^EC^LSN_HEK_)

Plasmids encoding ^EC^LSN_WT_ and ^EC^LSN_HEK_ were transformed into chemically competent *E. coli* BL21(DE3) cells (New England BioLabs). Cells were incubated with plasmid DNA on ice for 30 min, heat-shocked at 42 °C for 10 s, recovered in SOC medium for 1 h at 37 °C, and plated on LB agar containing kanamycin (50 µg/mL). Protein expression was performed using auto-induction medium based on modified ZYM-5052 medium. Starter cultures were grown in LB with antibiotics at 37 °C and diluted 1:200 into auto-induction medium supplemented with 0.5% (w/v) glycerol, 0.5% (w/v) glucose, and 0.2% (w/v) lactose. Expression cultures (2 L total) were incubated at 25 °C for 24 h with shaking. Cells were harvested by centrifugation at 4 °C. Cell pellets were resuspended in lysis buffer (20 mM HEPES [pH 7.0], 300 mM NaCl, 20 mM imidazole) and lysed by sonication. Lysates were treated with 0.01% (v/v) Benzonase, clarified by centrifugation, and applied to a 5 mL HisTrap HP column on an ÄKTA pure system. After washing with 40 mM imidazole, bound proteins were eluted using a stepwise imidazole gradient (200 and 800 mM). Fractions containing the target protein were pooled and analyzed by SDS–PAGE.

#### SDS–PAGE purity assessment

Lysin purity and integrity were evaluated by SDS–PAGE under reducing and non-reducing conditions. Samples (2.5 µg) were resolved on AnykD Criterion TGX precast gel (BioRad 5671125) and visualized by InstantBlue® Coomassie Protein Stain (Abcam, ab119211) staining. Purity was estimated by densitometric analysis using ImageQuantTL v10.2 software, and preparations exhibiting ≥95% purity were used for further analyses.

#### Size-exclusion chromatography (SEC-HPLC)

Size distribution and aggregation were assessed by size-exclusion chromatography using a TSKgel® G3000SWXL column (5 µm) coupled to an UltiMate™ 3000 HPLC system. Proteins were eluted in 20 mM HEPES, 150 mM NaCl, 300 mM Arginine (pH 7.0) buffer at a flow rate of 0.3 mL/min, and elution profiles were monitored at 280 nm. Monomer content was determined by peak area integration and was required to exceed 95%.

#### Intact mass analysis

Protein was analyzed without and with prior deglycosylation by incubation with recombinant PNGase F (Promega) overnight at 37 °C. Reduction was performed in 10 mM DTT / 3 M Guanidinium hydrochloride for 15 min at 65 °C before dilution to approximately 30 ng/µL with dH₂O. Sixty nanograms (2 µL) were loaded onto an XBridge Protein BEH C4 column (2.5 µm particle size, 2.1 mm × 150 mm; Waters) using a Vanquish™ Horizon UHPLC system (Thermo Scientific) operated at 50 °C. Solvent A consisted of 0.1% formic acid in water and solvent B of 100% acetonitrile containing 0.08% formic acid. Proteins were separated using a 6-min step gradient from 12 to 72% solvent B at a flow rate of 250 µL/min and analyzed on a Synapt G2-Si mass spectrometer coupled via a ZSpray ESI source (Waters). Data were acquired using MassLynx v4.2 (Waters) and deconvoluted with the MaxEnt1 algorithm to reconstruct the average neutral protein mass.

#### Nano Differential Scanning Fluorimetry (NanoDSF)

Thermal stability of purified proteins was assessed using a Prometheus NT.48 instrument (NanoTemper Technologies). Protein samples were prepared at a final concentration of 0.5 mg/mL in formulation buffer. Samples were thawed immediately prior to measurement. Standard capillaries (NanoTemper Technologies) were filled with approximately 10 µL of sample and loaded into the instrument. All measurements were performed in duplicates. Samples were heated from 20 °C to 95 °C at a constant ramp rate of 1.0 °C/min. Intrinsic protein fluorescence was monitored with excitation at 280 nm and emission recorded at 330 and 350 nm. Thermal unfolding transitions were determined from changes in the fluorescence emission ratio (F₃₅₀/F₃₃₀). In parallel, changes in light scattering were monitored as attenuation units to assess sample aggregation or turbidity during thermal unfolding.

#### Generation of the lysostaphin-reactive antibody

##### Antigen Preparation

*^EC^*LSN_HEK_ was expressed in *E. coli* and purified by Ni-NTA affinity chromatography as described above. Protein eluates were treated with HRV 3C protease (produced in-house, 1:100 protease:lysin molar ratio) while being dialyzed overnight against 300 sample volumes of final storage buffer (20 mM HEPES (pH 7.0), 150 mM NaCl). Purity (≥85%) was confirmed by SDS–PAGE followed by Coomassie staining. Aliquots were stored at –80 °C until immunization.

##### Rabbit Immunization

Polyclonal antibodies were raised by Genscript in New Zealand White rabbits under a 3-immunization protocol conducted in an AAALAC/OLAW-accredited facility. Pre-immune serum was collected before subcutaneous administration of antigen; two boosters were given at standard intervals; post-immune serum was collected after the final boost.

##### Affinity Purification and Cross-Adsorption

Pooled antisera (40 mL per rabbit) were subjected to antigen-affinity purification, yielding LSN–specific IgG in phosphate buffered saline (PBS [pH 7.4]) with 0.02% ProClin 300 and ≥99% purity by SDS–PAGE. To remove unwanted anti-IgG reactivity, antibodies were sequentially cross-adsorbed against immobilized human and mouse total IgG (≥85% purity). The final preparation yielded 9.31 mg total IgG at 0.776 mg/mL.

#### Western blot

Lysin integrity was assessed by SDS–PAGE followed by western blotting. Protein samples were resolved on AnykD Criterion™ TGX™ precast gels (Bio-Rad, 5671125) and were transferred onto 0.2 μm nitrocellulose membranes using the Trans-Blot® Turbo™ Transfer System (Bio-Rad, 1704271) applying the manufacturer’s Mixed Molecular Weight protocol. Membranes were blocked for at least 1 h at room temperature in 5% (w/v) skim milk prepared in Tris-buffered saline containing 0.05% (v/v) Tween-20 (TBS-T; Carl Roth, 1061.1). Blocked membranes were incubated overnight at 4 °C with anti-LSN primary antibody diluted 1:50 000 in 5% skim milk in TBS-T. After three washes, the membranes were incubated for 1 h at room temperature with horseradish peroxidase (HRP)–conjugated goat anti-rabbit IgG secondary antibody (Thermo Fisher Scientific, G-21234) diluted 1:10 000 in 5% skim milk in TBS-T. After three additional washes, signals were detected using Pierce™ ECL substrate (Thermo Fisher Scientific, 32106) and imaged on an ImageQuant™ 800 system (Cytiva). Densitometric analyses were performed using ImageQuantTL v10.2 software.

#### LC–MS/MS–based mapping of proteolytic cleavage sites

*Sample preparation for mass spectrometry analysis.* The Coomassie-stained gel band with an apparent molecular weight (Mw) of ∼50 kDa was excised from the gel and cut into 1 mm cubes, followed by destaining steps with a mixture of acetonitrile and 50 mM ammonium bicarbonate (ABC). The proteins were reduced using 10 mM DTT in 50 mM ABC and alkylated with 50 mM iodoacetamide in 50 mM ABC. Digestion was carried out with trypsin (Promega; Trypsin Gold, Mass Spectrometry Grade) in 50mM ABC at 37°C overnight. Formic acid was used to stop digestion and for peptide extraction.

#### Liquid chromatography–tandem mass spectrometry (LC–MS/MS)

LC-MS/MS analysis was performed on a Vanquish Neo UHPLC system (Thermo Scientific) coupled to an Orbitrap Exploris 480 mass spectrometer (Thermo Scientific). The system was equipped with a Nanospray Flex ion source (Thermo Scientific). Peptides were loaded onto a trap column (PepMap Neo C18 5mm × 300 µm, 5 μm particle size, Thermo Scientific) using 0.1% trifluoroacetic acid as mobile phase, and separated on an analytical column (Acclaim PepMap 100 C18 HPLC Column, 50 cm × 75 µm, 2 μm particle size, Thermo Scientific) applying a linear gradient starting with a mobile phase of 98% solvent A (0.1% formic acid) and 2% solvent B (80% acetonitrile, 0.08% formic acid), increasing to 35% solvent B over 60 min at a flow rate of 230 nl/min. The analytical column was heated to 30°C. The mass spectrometer was operated in data-dependent acquisition mode, with 2 s MS1 cycle time. Survey scans were acquired from 375-2000 m/z with lock mass enabled, normalized AGC target of 300%, resolution of 60,000. The most intense precursor ions (charge states +2 to +6) were selected for fragmentation using an isolation window of 1.4 m/z. Selected ions were analyzed with a maximum fill time of 200 ms, normalized automatic gain control target of 200%, and resolution of 30,000 after higher-energy collisional dissociation fragmentation with normalized collision energy of 30%. Monoisotopic precursor selection was set to “peptide” mode, the intensity threshold to 2.5 × 10^4^, and selected precursors were dynamically excluded for 20 seconds with isotope exclusion enabled.

#### FragPipe

MS raw data were analyzed with FragPipe (24.0), using MSFragger (4.4.1) (Kong et al., 2017), IonQuant (1.11.20) (Yu et al., 2021), and Philosopher (5.1.1) (da Veiga Leprevost et al., 2020). The default FragPipe workflow for label free quantification was used. Cleavage specificity was set to Trypsin/P, with 3 missed cleavages allowed. Methionine oxidation and protein N-terminal acetylation as well as trioxidation and carbamidomethylation of cysteine were specified as variable modifications. MS2 spectra were searched against the Fc-L2a-CHAP_opt_var_-LSN_HEK_ sequence with *H. sapiens* 1 protein per gene reference proteome from Uniprot (Proteome ID: UP000005640, release 2026.01), concatenated with a database of common laboratory contaminants (release 2026.01 https://github.com/maxperutzlabs-ms/perutz-ms-contaminants).

### Bacterial strains, culture conditions and substrate generation

Staphylococci were streaked for single colonies from cryo vials onto Columbia Agar with 5% sheep blood (BD, 221263) and incubated statically overnight at 37 °C. Plates were stored at 4 °C for up to 2 weeks. Several colonies were used to inoculate 10 mL tryptic soy broth (TSB, Carl Roth) cultures in 50 mL Falcon tubes and incubated overnight at 37 °C with shaking at 220 rpm. If mid-log cultures were required, the overnight cultures were diluted in TSB to OD_600_=0.05 and grown at 37 °C with shaking at 240 rpm for 2-3 h to OD₆₀₀ ≈ 0.7–1.5. The following strains were used for this study: *Staphylococcus aureus* ATCC 43300 (ATCC), BAA-1717 (ATCC), NCTC 8178 (NCTC), *Staphylococcus epidermidis* DSM3269 (DSMZ), *S. epidermidis* RP62A (ATCC), *Staphylococcus lugdunensis* DSM4804 (DSMZ), *Staphylococcus haemolyticus* DSM20263 (DSMZ) and *Staphylococcus haemolyticus* DSM20265 (DSMZ).

#### Generation of peptidoglycan substrate for western blot

A single colony of *S. aureus* ATCC43300 from a streak out plate was inoculated in 10 mL of BHI and grown overnight at 37°C, shaking at 250 rpm. After 16 hours of growth a mid-log culture was set up by diluting the overnight culture 1:100 in 200 mL fresh BHI in a sterile 1 l bottle and incubated approximately for 3 hours to OD_600_ = 3. The culture was autoclaved and washed twice with sterile water (centrifugation 8000 rpm for 10 min) followed by freezing of the pellet at -20°C until further use.

### Functional and biophysical characterization

#### Growth inhibition assays for the determination of minimal inhibitory concentrations (MICs)

MIC was determined using a microbroth dilution method following the CLSI Standard M100-Ed35 Appendix H2 for antimicrobial susceptibility testing (CLSI, 2026). 2-fold dilution series of lysins were prepared as 5× stock solutions in cation adjusted MHB (caMHB, Sigma Aldrich) supplemented with 20% heat-inactivated horse serum (HS; Thermo Fisher Scientific, 26050070). Lysins were tested at 0.25-32 µg/mL. Mid-log bacterial cells were adjusted to OD_600_=0.001 in caMHB with 20% HS, to reach a final inoculum of 5×10^5^ colony forming units per mL (CFU/mL). Reactions were set up in triplicate in 96-well U-bottom plates by mixing 20 µL of lysin solution with 80 µL bacterial suspension. Plates were statically incubated at 37 °C for 17-19 h. MIC was defined as the lowest concentration of lysin that still prevented growth by visual inspection and determined for each replicate.

#### Time kill assay (TKA)

Mid-exponential-phase bacteria (see above) were harvested, washed twice in Dulbecco’s Phosphate Buffered Saline (DPBS) containing Ca²⁺/Mg²⁺ (Thermo Fisher Scientific, 14040174), and resuspended to OD₆₀₀ = 0.125 in human serum (HuS; Innovative Research, ISER500ML) or DPBS containing Ca²⁺/Mg²⁺. If instead autoclaved cell pellets were used, autoclaved bacterial pellets were thawed, resuspended in DPBS supplemented with Ca²⁺/Mg²⁺, and adjusted to an OD₆₀₀ of 0.5 in HuS. Lysins were diluted to 5 μM in HEPES buffer (20 mM HEPES [pH 7], 150 mM NaCl) or MES buffer (50 mM MES [pH 6.5], 300 mM NaCl, 1 mM CaCl2, 2 mM DTT, 5% glycerol, and 50 mM arginine), further diluted to working solutions in HEPES buffer supplemented with 2.5 % (w/v) bovine serum albumin (BSA). Reactions were set up in sterile 96-well flat-bottom plates by mixing 20 μl lysin with 80 μl of the OD-adjusted bacterial suspension. Plates were incubated statically at 37 °C for 1 h, 3 h, 5 h and 24 h. Aliquots were collected for quantitative plating, SDS-PAGE and western blot analysis. Quantitative spotting was performed by five 10-fold serial dilutions in DPBS and spotting 2 µL of the dilutions onto Trypton-Soja-Agar (TSA) plates (BD, 221803). After overnight incubation at 37 °C, CFU/mL were determined. The log10 reduction in CFU/mL was calculated relative to the buffer-treated growth control.

#### Flat Bottom Adherent Biofilm Assay (FBABA)

Overnight cultures were diluted in caMHB supplemented with 10 % pooled, sterile-filtered and dialyzed (1× DPBS using 10 kDa dialysis membrane) human plasma supplemented with natrium citrate (caMHB–10% HuP) to OD₆₀₀ = 0.1 (∼1 × 10⁸ CFU/mL). Bacterial input concentrations were confirmed by serial dilution and quantitative spotting (see above). Aliquots (100 µL/well) were transferred into tissue culture–treated 96-well plates and incubated statically at 37 °C for 24 h to allow biofilm formation. Planktonic cells were removed by blotting. Biofilms were gently washed three times with 1× DPBS and treated with 125 µL antimicrobial solutions prepared in caMHB supplemented with 50% human plasma (caMHB–50% HuP) for 24 hours at 37 °C. Biofilms were washed three time with 1× DPBS. The biofilm was dislodged using 100 µL TrypLE (Thermo Fischer, 12604013) for 20 min at 37 °C and fully resuspended upon the addition of 100 µL 1× DPBS. Viable bacteria were quantified by ten-fold serial dilution and spot plating on TSA. Biofilm reduction was calculated as CFU/mL and expressed as log10 CFU reduction relative to untreated controls.

#### SEC-SAXS

Small-angle X-ray scattering (SAXS) data were acquired by BIOSAXS GmbH at the P12 beamline of the PETRA III synchrotron (DESY, Hamburg, Germany). The undulator (gap 20.95 mm) and monochromator were tuned to an incident energy of 10 keV. Beamline calibration of the angular axis was performed using silver behenate. Prior to data collection, beamline performance was verified following the P12 standard operating procedures, including measurements of empty capillaries, Milli-Q water, BSA, and the corresponding HEPES buffer.

Technical Specifications of P12 beamline used for the analysis.

|  |  |
| --- | --- |
| Source | Petra III U29 undulator, P12 BioSAXS beamline |
| Detector | hybrid photon detector Pilatus 6M |
| Energy range | 10 keV |
| Maximum flux at sample | $1 \times 10^{12}$ photons per second |
| Beam size at sample | 250 μm × 100 μm |
| s-range | 0.03 – 7.25 nm<sup>-1</sup> |
| Wavelength | 0.124 nm |
| Sample-to-Detector Distance | 3 m |
| Number of bunches | 40 |
| PETRA III ring current | 100 mA |
| Axis Calibration Standard | silver behenate |
| Beamstop position | off-set |

Samples were subjected to inline size-exclusion chromatography–coupled SAXS (SEC–SAXS) using an Agilent Bioinert Infinity 1260 HPLC system connected to a Cytiva Superdex 200 Increase 5/150 column. The column was equilibrated with 10 CV of SEC buffer (20 mM HEPES, 150 mM NaCl, pH 7.0) and linked directly to the SAXS flow cell. For each SEC–SAXS run, 25 μL of protein sample at 5 mg/mL were injected. A total of 1,800 scattering frames were recorded during elution of the protein peak and subjected to automated radial averaging as previously described (Blanchet et al., 2015; Hajizadeh et al., 2018). Data processing and extraction of global scattering parameters were performed using CHROMIXS (Panjkovich and Svergun, 2017) and PRIMUS (Konarev et al., 2003), including calculation of real-space distance distribution functions. Ab initio shape reconstruction was carried out as described by Svergun et al, 1999 (data not shown), yielding overall shapes of Fc-L1-CHAP_opt_-LSN_HEK_ with the following parameters:

|  |  |
| --- | --- |
| Rg1 (nm) | 7.3 ± 0.1 |
| Rg2 (nm) | 8.8 ± 0.1 |
| Dmax (nm) | 40 ± 3 |
| Vdam (nm<sup>3</sup>) | 230 ± 15 |
| MWdam (kDa) | 120 ± 10 |

with R_g1_ being the radius of gyration determined from the Guinier approximation, R_g2_ the radius of gyration determined from the distance distribution function computed by the program GNOM (Svergun, 1992), D_max_ the maximum particle size assessed from the p(r) function in GNOM, V_dam_ the volume of a typical ab initio model reconstructed by the program DAMMIN (Svergun, 1999) and MW_dam_, the approximate molecular weight assessed from the typical DAMMIN model.

Rigid-body modelling of Fc-L1-CHAP_opt_-LSN_HEK_ was performed with CORAL (Petoukhov et al., 2012). The protein was represented as a dimer comprising the IgG1 Fc domain (PDB: 6eaq), a helical linker derived from PDB:6jfv, the CHAP domain (PDB: 11CI) and LSN (PDB: 4LXC), with the latter two connected by a short GG linker. The dimeric nature of Fc was placed at the origin and oriented with the principal axes along X, Y and Z. Multiple CORAL runs were conducted on the Fc-L1-CHAP_opt_-LSN_HEK_ data and the restored symmetric dimers displayed an overall similar appearance yielding a good fit to the data. Two different organizations of the monomers in the dimer were observed, namely trans-dimers (as shown in Figure 8A) and cis-dimers (data not shown), with the CHAP-LSN arms located either on the opposite or same site, respectively, of the fused Fc domain. Both isoforms displayed otherwise similar conformations and dimensions.

#### Lytic assay with spiked-in LSN

A master stock of 10 µM Fc-L1-CHAP_opt_-LSN_HEK_ was prepared in assay buffer (50 mM MES [pH 6.5], 300 mM NaCl, 1 mM CaCl_2_, 2 mM DTT, 5% [v/v] glycerol, 50 mM arginine and 2.5% [w/v] BSA). LSN_HEKa_ was prepared as a 1 µM master stock in the same buffer and serially diluted two-fold. Log-phase *S. aureus* ATCC 43300 cells were harvested (4000 × g, 5 minutes), washed twice with 1× DPBS and resuspended in either 1× DPBS or HuS to OD_600_ = 1.25. Lytic assays were performed in 96-well flat-bottom plates by mixing 120 μl of bacterial suspension with 15 µl Fc-L1-CHAP_opt_-LSN_HEK_ (10 µM) and 15 µl LSN_HEKa_ (500 - 62.5 nM). Control samples contained the single agents adjusted with assay buffer or the buffer alone. Samples were statically incubated for 0, 1 or 3 hours at 37°C. Samples were collected for quantitative spotting (see above) and western blot analysis. The LSN band intensity (∼27 kDa) were using ImageQuant™TL v10.2. A standard curve correlating band intensity with LSN concentration was generated from 0-h samples and used to calculate LSN concentrations at later time points. Killing efficacy was expressed as log10 CFU/mL reduction and plotted against calculated LSN concentrations.

### In vivo studies

Animal studies were performed at Evotec (UK) under UK Home Office Licenses and with local ethical committee clearance. All experiments were performed in dedicated BSL2 facilities. SPF (specific pathogen free) CD1 mice used in this work were supplied by Charles River Laboratories (Margate, UK). After arrival, mice were acclimatized for 7 days and housed in IVC cages at 22 ± 1 °C, 60 % relative humidity, maximum background noise of 56 dB and under 12-hour light/dark cycles. They had free access to food, sterile water, and were provided with aspen chip bedding. When needed, animals were supplemented with wet food mush and HydroGel for additional hydration.

#### Inoculum preparation and quantitative burden

Infections were performed with freshly prepared *S. aureus* (ATCC 43300 or NCTC 8178) grown to logarithmic phase. The target inoculum was confirmed by quantitative plating on mannitol salt agar (MSA) plates. Mice were euthanized by terminal cardiac bleed under isoflurane anesthesia and death confirmed by cervical dislocation. Tissue samples were homogenized in PBS using Precellys homogenizer and homogenates were serially diluted 1:10 in sterile PBS, followed by quantitative plating on MSA plates. Plates were incubated at 37 °C for 18-24 h and bacterial colonies enumerated.

#### Mouse ex vivo catheter infection model

Efficacy of Fc-L1-CHAP_opt_-LSN_HEKa_ against biofilms was tested in a mouse model of *S. aureus* ATCC 43300 foreign body (catheter) infection. Five days before bacterial challenge CD1 mice have undergone surgery where a FEP (Fluorinated Ethylene Propylene) catheter fragment (dimension L∼10 mm × dia. 2.1 mm, BD Angiocath) was subcutaneously implanted on the flank. Mice were infected by transcutaneous injection of 100 µL containing 5×10^6^ CFU *S. aureus* ATCC 43300, directly into the catheter. Biofilms were allowed to develop *in vivo* for 3 days, after what the catheters were explanted and the formed biofilms were treated with vehicle or Fc-L1-CHAP_opt_-LSN_HEKa_, at a concentration of 50 µg/mL or 100 µg/mL in caMH broth with 50 % plasma, for 24 h at 37 °C. Biofilms were dislodged by sonication and quantitative catheter biofilm burden was determined through quantitative plating techniques.

#### Mouse IV sepsis model

Mice were infected with a 100 µL injection in the lateral tail vein of 5×10^7^ *S. aureus* NCTC 8178. Body weights were recorded and weight loss relative to day 0 calculated over the course of the study (ethical limit >20% weight loss). The clinical condition of mice was observed and scored regularly throughout the study to monitor animal welfare. Recombinant protein lysins were administered IV twice, at 1 hour post infection (hpi) and 25 hpi. Fc-L1-CHAP_opt_-LSN_HEK_ was given at dose of 500 µg/mouse, while Fc-L2-CHAP_opt_-LSN_HEK_ was given at two dose levels, 100 µg/mouse and 500 µg/mouse. Vancomycin treatment (25 mg/kg, IV, q12 h from 1 hpi) was used as comparator. Quantitative bacterial burden in both kidneys together was determined at 72 hpi (pre-treatment at 1 hpi).

## Results

### Construct design

We initially generated half-life extended aglycosylated LSN by fusing it either N-terminally or C-terminally to the human IgG1 Fc with a helical linker containing 8 repeats (L1), leading to LSN_HEKa_-L1-Fc or Fc-L1-LSN_HEKa_, respectively (Supplementary Figure 1). The fusions retained high bacteriolytic activity (low MIC) against multiple *S. aureus* strains, with LSN_HEKa_-L1-Fc being less impaired by the Fc-fusion than Fc-L1-LSN_HEKa_ (Table 1). Yet, unlike free LSN, the half-life extended constructs were less active or even inactive against coagulase negative staphylococcal (CNS) strains *S. epidermidis*, *S. haemolyticus* or *S. lugdunensis* (Table 2).

**Table 1:**
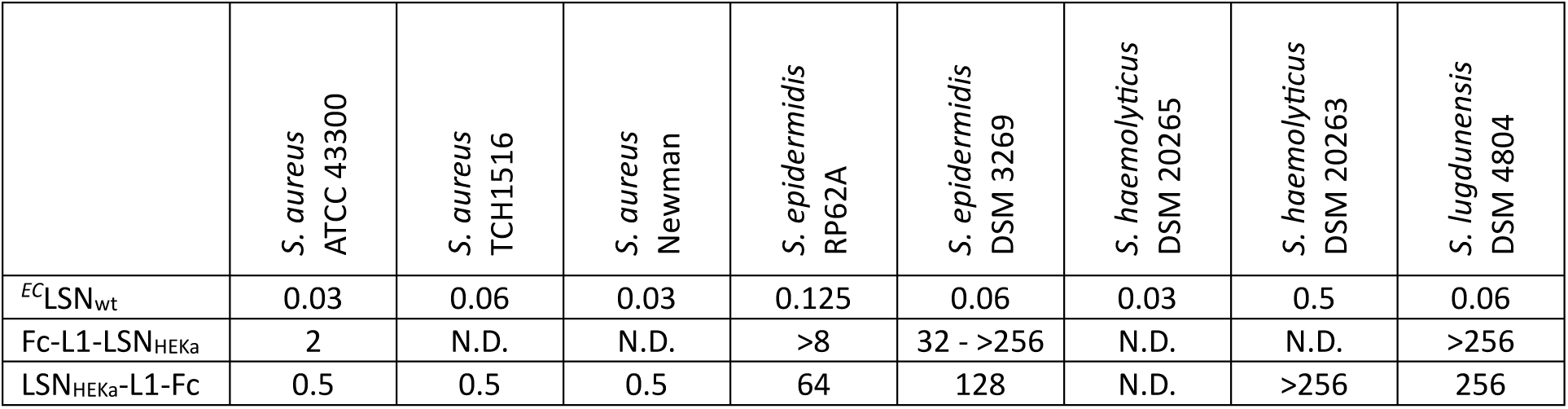
Minimal inhibitory concentration of LSN and its Fc fusions for *S. aureus* and CNS strains. The MICs in this table are shown in µg/mL. N.D. indicates that the MIC was not determined.

**Table 2:** Median yields and minimal inhibitory concentrations of Fc-CHAP-LSN variants. The data is compiled over purifications from multiple batches and formulation buffers. The MICs are reported in µg/mL for *S. aureus* and three CNS strains. Data from single experiment is marked with “#”, N.D. indicates the data was not determined.

| Lysin | Yield ($\mu\text{g/mL}$) | MIC<br>*S. aureus*<br>ATCC 43300 | MIC<br>*S. lugdunensis*<br>DSM 4804 | MIC<br>*S. epidermidis*<br>DSM3269 | MIC<br>*S. epidermidis*<br>RP62A |
| --- | --- | --- | --- | --- | --- |
| Fc-L1-CHAP-LSN<sub>HEK</sub> | 9.8 | 2 | 32 | 39 | 4 |
| Fc-L1-CHAP<sub>opt</sub>-LSN<sub>HEK</sub> | 45.2 | 1 | 2 | $\geq 128$ | N.D. |
| Fc-L1-CHAP<sub>opt</sub>-LSN<sub>HEKa</sub> | 52.8 | 2 | 38 | 64 | N.D. |
| Fc-L2-CHAP<sub>opt</sub>-LSN<sub>HEK</sub> | 49.4 | 1 | 1<sup>#</sup> | N.D. | N.D. |
| Fc-L1-CHAP*-LSN<sub>HEK</sub> | 61.0 | 8 | >256 | >256 | >128 |
| Fc-L1-CHAP-LSN*<sub>HEK</sub> | 11.7 | >64 | N.D. | N.D. | N.D. |

To extend the substrate profile of the lysin, we generated fusions with additional enzymatic domains. As a first attempt, we fused different CHAPs to the LSN sequence, connected by a Gly-Gly linker. We postulated that the fusion of a CHAP with LSN (a Zn^2+^-dependent peptidase) could yield a lysin with two different modes of action and potentially overcome the known resistance formation observed for LSN (Becker et al., 2016). The introduction of an engineered CHAP domain (the CHAP fragment from LysM-CHAP L3 in Badarau et al., 2026 led to lysin Fc-L1-CHAP-LSN_HEK_ with a MIC of 2 µg/mL on *S. aureus* and restored activity on CNS strains (Table 2). The MIC of Fc-L1-CHAP-LSN_HEK_ on *S. lugdunensis* strain DSM 4804 was 32 µg/mL, eight times lower than for LSN_HEKa_-L1-Fc, validating the hypothesis that the addition of the CHAP to LSN increases lysin strain coverage. Interestingly, when the CHAP catalytic cysteine was mutated to serine, yielding enzymatically inactive CHAP (Fc-L1-CHAP*-LSN_HEK_), the activity on CNS strains was lost and the activity on *S. aureus* was decreased, confirming that both catalytic domains contribute to the bactericidal activity of the lysin.

However, Fc-L1-CHAP-LSN_HEK_ had ∼ 7-fold lower yields than Fc-L1-LSN_HEKa_ (data not shown), and we set out to engineer the CHAP domain by introducing selected mutations, to increase expression and yield. One such mutation, namely tyrosine 46 to aspartate (Y46D) identified previously (Badarau et al., 2026), yielding Fc-L1-CHAP_opt_-LSN_HEK_ and Fc-L1-CHAP_opt_-LSN_HEKa_ chimeric lysins, resulted in a 5-fold improvement in yield compared with Fc-L1-CHAP-LSN_HEK_, without compromising activity (Table 2). Further optimization steps included elongating the helical linker between Fc and CHAP-LSN from 8 to 14 helical repeats and appending a 12-amino-acid segment derived from serum stable LysM-CHAP linkers (PKETVSLNEESN, L2; or KPETVSGNEKSN, L2a) (Supplementary Figure 1) (Badarau et al., 2026). Several formulation buffers were tested during optimization, revealing that Fc-CHAP-LSN constructs were most stable in MES buffer at pH 6.5, as judged from melting temperatures (T_m_) and final purification yields. The engineering campaigns led to increases in T_m_ of ∼3°C (Fc-L1-CHAP-LSN_HEK_ compared with Fc-L2a-CHAP_opt_-LSN_HEK_) and in final yield by more than 6-fold from Fc-L1-CHAP-LSN_HEK_ to Fc-L2-CHAP_opt_-LSN_HEK_ (Table S1).

To assess the activity of Fc-CHAP-LSN constructs in a biofilm context, a series of experiments was performed comparing Fc-CHAP-LSN constructs with LSN_HEKa_-L1-Fc as well as to standard of care (SOC) antibiotics. As had already been determined in the MIC experiments, Fc-CHAP-LSN constructs outperformed Fc-tagged LSN and SOC antibiotics in reducing *S. aureus* biofilms: Fc-L1-CHAP-LSN_HEK_ at a dose of 12.5 µg/mL led to ∼3 log10 reduction in bacterial count, whereas LSN_HEKa_-L1-Fc only reduced the bacterial burden by 2 logs (Figure 1). Interestingly and in contrast to Fc-L1-LSN_HEKa_, no dose dependency was seen for Fc-L1-CHAP-LSN_HEK_. Compared with the standard-of-care antibiotic used at concentrations approximating the maximum plasma concentration (C_max_) in humans, both lysins showed superior antibiofilm activity.

**Figure 1:**
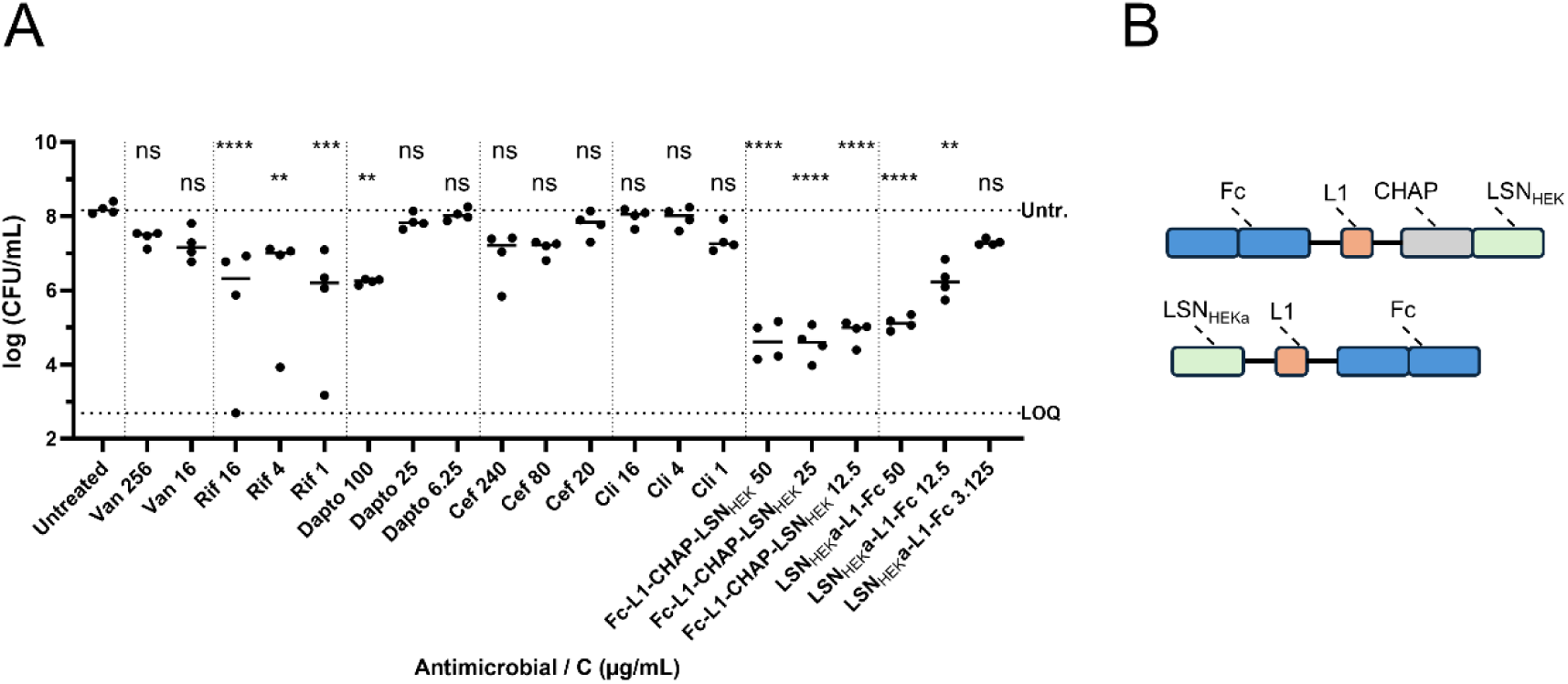
(A) Efficacy of Fc-CHAP-LSN constructs against *in vitro* formed biofilms. Bacterial quantification of 24 h-cultivated biofilms of *S. aureus* ATCC 43300 after 24 h exposure to SOC antibiotics or lysins. Concentrations indicated in µg/mL. Statistical analysis was performed using Graphpad’s one-way ANOVA with Dunnett’s correction for multiple comparisons; ****: P<0.0001; ***: P=0.001; **: P<0.01 ns: not significant (B) Graphical representation of the tested lysins in this assay. Cli: clindamycin; Dapto: daptomycin; LOQ: limit of quantification at 500 CFU/mL; Rif: rifampicin; Untr.: untreated control; Van: vancomycin;

To evaluate anti-biofilm activity in an *in vivo*-like setting, we ran an *ex vivo* catheter foreign body infection model (Heim et al., 2014). Briefly, Fluorinated Ethylene Propylene (FEP) catheters were implanted into CD1 mice followed by infection with *S. aureus* ATCC43300 4 days post implantation. Biofilm was allowed to establish for 3 days, after which the catheters were explanted and treated *in vitro* with Fc-L1-CHAP_opt_-LSN_HEKa_ at 50 or 100 µg/mL or SOC comparator rifampicin at 10 µg/mL (Figure 2A). The catheter bacterial biofilm burden was determined 24 h after single treatment. As shown Figure 2B, *ex vivo* treatment with Fc-L1-CHAP_opt_-LSN_HEKa_ at 50 µg/mL and 100 µg/mL resulted in a significant 4.0 (P<0.0001) and 3.1 (P=0.0105) log10 CFU/catheter reduction in biofilm bacterial burden, respectively, compared with the vehicle-treated group. In contrast, bacterial burden reduction with rifampicin treatment was 2.0 log10 CFU/catheter vs vehicle (P=0.1441). Following successful *ex vivo* biofilm treatment, we aimed at assessing *in vivo* efficacy in a 72 h murine intravenous (IV) sepsis model to better mimic the clinical situation (Badarau et al., 2026). Treatment was started 1 h post infection, with 2 doses of 500 or 100 µg/mouse given 24 h apart for the lysins and comparator vancomycin dosed every 12 h (q12 h) (Figure 2C). Vancomycin as well as Fc-L2-CHAP_opt_-LSN_HEK_ at 100 µg/mL led to approximately 5.0 and 4.0 log10 reduction respectively (P<0.0001) (Figure 2D). Fc-L2-CHAP_opt_-LSN_HEK_ as well as Fc-L1-CHAP_opt_-LSN_HEK_ at 500 µg/mouse led to a similar 5.0 log10 reduction (P=0.0385 and P=0.0136 respectively). Interestingly, in the *ex vivo* catheter model as well as in the murine sepsis model, all Fc-CHAP-LSN constructs tested were highly efficacious, yet no dose dependency was observed in either model (Figure 2B and Figure 2D), which prompted us to investigate the mode of action of this lysin architecture.

**Figure 2:**
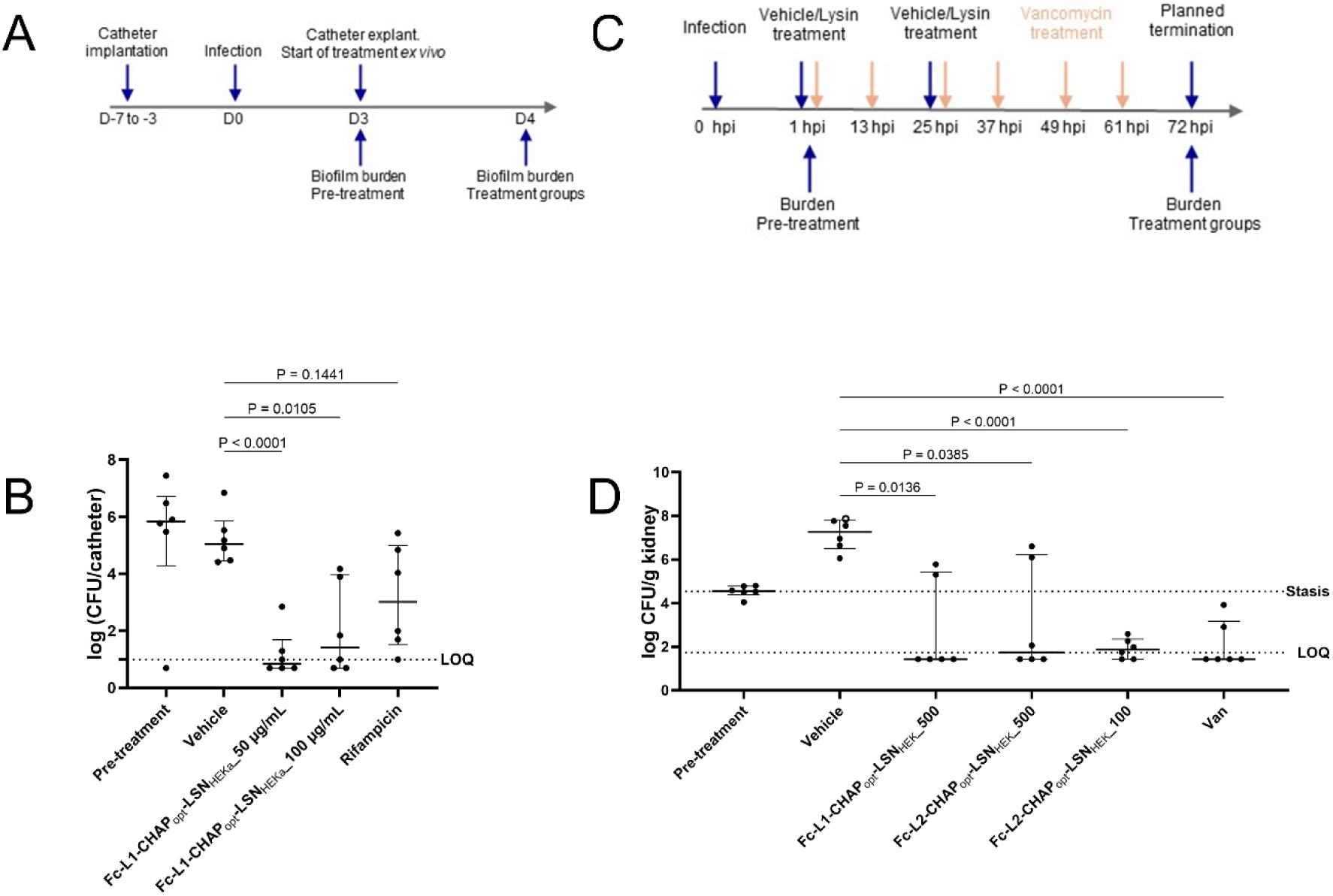
Efficacy of Fc-CHAP-LSN against *in vivo* formed biofilms and in sepsis model. (A) Catheters with *in vivo* formed biofilms of *S. aureus* ATCC 43300 were explanted from CD1 mice 3 dpi and treated with either 50 µg/mL or 100 µg/mL Fc-L1-CHAP_opt_-LSN_HEKa_ for 24 h at 37 °C. Rifampicin (10 µg/mL) was used as comparator. (B) Datapoints represent the biofilm bacterial burden in each catheter (n=6). Horizontal bars indicate the median burden for each group. When burden is below LOQ, half a value of LOQ is assigned for graphical and statistical purposes. Brown Forsythe ANOVA with Welch’s correction (GraphPad Prism) was used to calculate statistical significance against vehicle. dpi: days post infection; LOQ: limit of quantification at 10 CFU/catheter. (C) Lysin efficacy in mouse IV sepsis: Immunocompetent male CD1 mice were challenged with 3.23 × 10^7^ CFU/mouse of S. aureus NCTC 8178, IV. Lysins were administered IV, twice (1 hpi and 25 hpi) at dose of either 100 µg/mouse or 500 µg/mouse. Vancomycin (Van) was dosed at 25 mg/kg, IV, q12 h, starting from 1 hpi. (D) Points on the graph represent CFU counts in individual mice (n=6), with horizontal lines indicating median CFU burden and interquartile range. Open circles indicate animals that needed to be terminated before planned endpoint due to clinical symptoms. Brown-Forsythe ANOVA with Welch’s correction was used to calculate the significance compared to vehicle group on log transformed data, with P values indicated in the graph. hpi: hours post infection; CFU: colony forming units; LOQ: limit of quantification at 54 CFU/g; For graphical and statistical purposes, values under LOQ were assigned half of the LOQ value.

### Mode of action

During test expressions of Fc-CHAP-LSN constructs composed of different CHAP domains and linkers, we consistently detected the presence of defined fragments of these constructs corresponding to the molecular weights of the CHAP-LSN module cleaved at different regions of the CHAP domain (33 and 37 kDa) and the LSN part (27kDa) (Figure 3).

**Figure 3:**
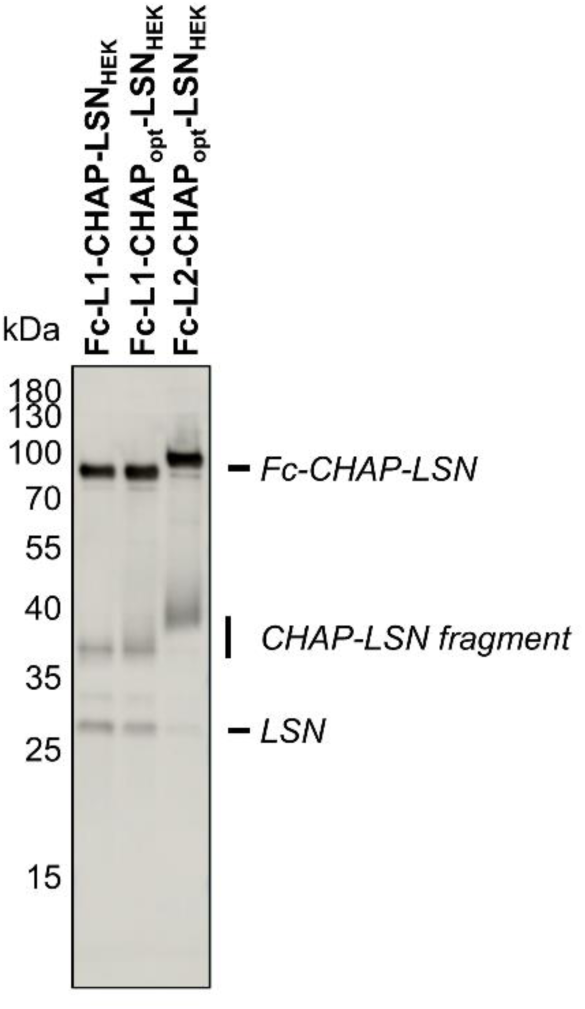
Fragmentation of Fc-CHAP-LSN after secretion from Expi293 cells. Variants Fc-L1-CHAP-LSN_HEK_, Fc-L1-CHAP_opt_-LSN_HEK_ and Fc-L2-CHAP_opt_-LSN_HEK_ were transfected into Expi293 cells. Supernatants were harvested and analyzed by SDS-PAGE followed by Western Blot. LSN containing species were detected using a LSN-directed antibody. Fragments corresponding to LSN, fragmented CHAP-LSN and full-length Fc-CHAP-LSN are indicated. Molecular weights in kilo Dalton (kDa) are indicated.

Mutation of the active site residues in either the CHAP domain, LSN, or both simultaneously significantly reduced the presence of these fragments, indicating that Fc-CHAP-LSN fragmentation is driven by the enzymatic activities of either the CHAP- or the PepM23 domain or both (Supplementary Figure 2A). This was unexpected for the CHAP domain, which acts specifically on the D-Ala-Gly bond only found in cell wall stem peptides (Frankel et al., 2011; Frankel and Schneewind, 2012). In contrast, PepM23 is known to cleave peptide bonds at Gly-Gly linkages, even if the target sequence is not part of the stem peptide (Grishin et al., 2020; Antenucci et al., 2024). Alanine scanning mutagenesis of extended glycine stretches in the CHAP domain (i.e. at positions CHAP_opt-85GGG87_ and CHAP_opt-115GGG117_) and lysostaphin (LSN_HEK-62GGG64_) indeed resulted in changes in the cleavage patterns, consistent with hydrolysis within the two poly-Gly regions in the CHAP domain (Supplementary Figure 2B). The LSN variants with mutations in the LSN_HEK-62GGG64_ loop were the only ones where the LSN (27 kDa) band disappeared and showed a strong reduction in both bactericidal activity and fragmentation in supernatants, suggesting that supernatant activity of Fc-CHAP-LSN is at least in part driven by the activity of the released LSN (Supplementary Figure 2C).

As a next step, Fc-L1-CHAP_opt_-LSN_HEK_ was purified to homogeneity from supernatants, which also removed any detectable traces of the CHAP-LSN and LSN fragments seen in supernatants (Supplementary Figure 3A). When tested for activity in TKAs against *S. aureus* ATCC 43300 in human serum, Fc-L1-CHAP_opt_-LSN_HEK_ showed a log10 CFU reduction of ∼1.5 relative to the growth control, consistent with other optimized and Fc-tagged lysin architectures (Badarau et al., 2026) (Figure 4A). Unexpectedly - and despite the fact that free LSN and CHAP-LSN had been removed during the purification of Fc-L1-CHAP_opt_-LSN_HEK_ - analysis of the TKA reactions by SDS-PAGE followed by western blotting indicated the presence of a significant amount of free LSN (∼0.0125 µg/mL) in the reaction – see below (Figure 4A, Supplementary Figure 3B), suggesting that release of LSN from Fc-CHAP-LSN had occurred during the reaction.

**Figure 4:**
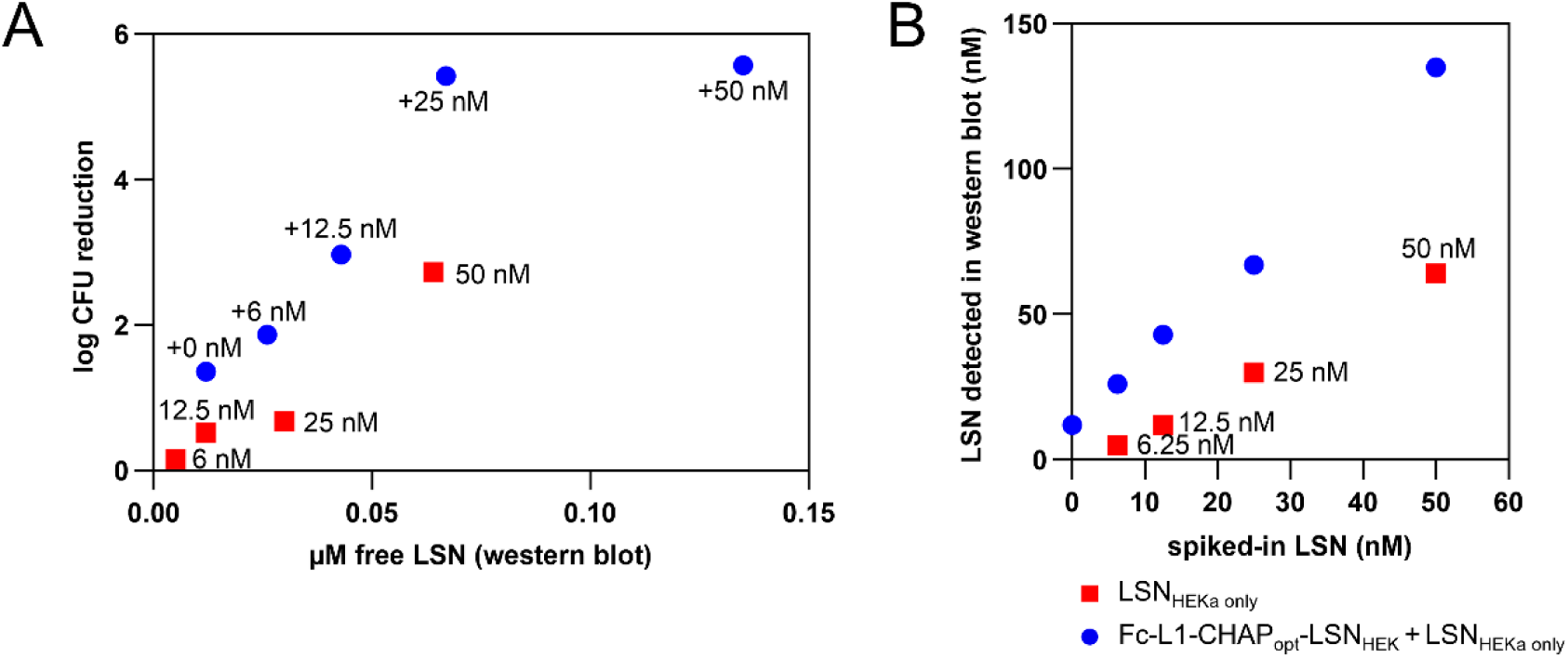
LSN-induced LSN-release from Fc-CHAP-LSN. (a) Purified LSN_HEKa_ was incubated at increasing concentrations in 80% human serum and in presence of 10^8^ CFU/mL *S. aureus* ATCC 43300 with (circles) or without (boxes) a fixed concentration of Fc-L1-CHAP_opt_-LSN_HEK_ (1 µM) for 3h at 37°C. Samples were analyzed by SDS-PAGE followed by Western Blotting to quantify the amounts of free LSN in each reaction (x-axis). The remainder of the reaction was subjected to quantitative plating to determine the log10 CFU reduction relative to a growth control lacking lysins (y-axis). (B) Concentrations of spiked-in purified LSN are plotted against free LSN detected by Western blotting in the experiment described in panel (A).

To quantify the impact of free LSN on the log10 CFU reductions in TKAs, we next titrated recombinant LSN produced in *E. coli* (^EC^LSN_WT_) into the reactions while maintaining constant levels of Fc-L1-CHAP_opt_-LSN_HEK_ (Figure 4A). We observed an initially linear increase in log10 CFU reduction with increasing levels of recombinant ^EC^LSN_WT_ which plateaued above ∼0.025 µM, where full bacterial eradication was observed. Interestingly, bactericidal activities of reactions in which Fc-L1-CHAP_opt_-LSN_HEK_ had been supplemented with recombinant LSN were drastically higher compared with reactions in which Fc-L1-CHAP_opt_-LSN_HEK_ or recombinant ^EC^LSN_WT_ had been added alone (Figure 4B).

Analysis of protein species by SDS-PAGE/western blot after TKA reactions showed that the levels of free LSN and CHAP-LSN fragments had increased with the addition of recombinant ^EC^LSN_WT_ in a dose dependent manner (Figure 4B). Interestingly, the concentrations of released LSN were up to three-fold higher compared with the amounts of recombinant LSN which had been added at the beginning of the experiment, suggesting that the presence of LSN in the reaction had promoted further release of LSN (and CHAP-LSN) from Fc-L1-CHAP_opt_-LSN_HEK_ (Figure 4B).

We next characterized how LSN-release is affected by the presence of human serum and target cells. LSN was released from Fc-L1-CHAP_opt_-LSN_HEK_ in presence of *S. aureus* cells both in human serum and DPBS in a cell density dependent manner, though the reaction was faster in human serum (∼ 1h) vs PBS (∼ 24 h) (Figure 5A, B, Supplementary Figure 4). In contrast, in the absence of *S. aureus* cells there was no or only minimal LSN release (Figure 5A, B). On the other hand, formation of the CHAP-LSN fragment required only the presence of human serum, with the same band intensities detected in the presence and absence of cells (Figure 5B).

**Figure 5:**
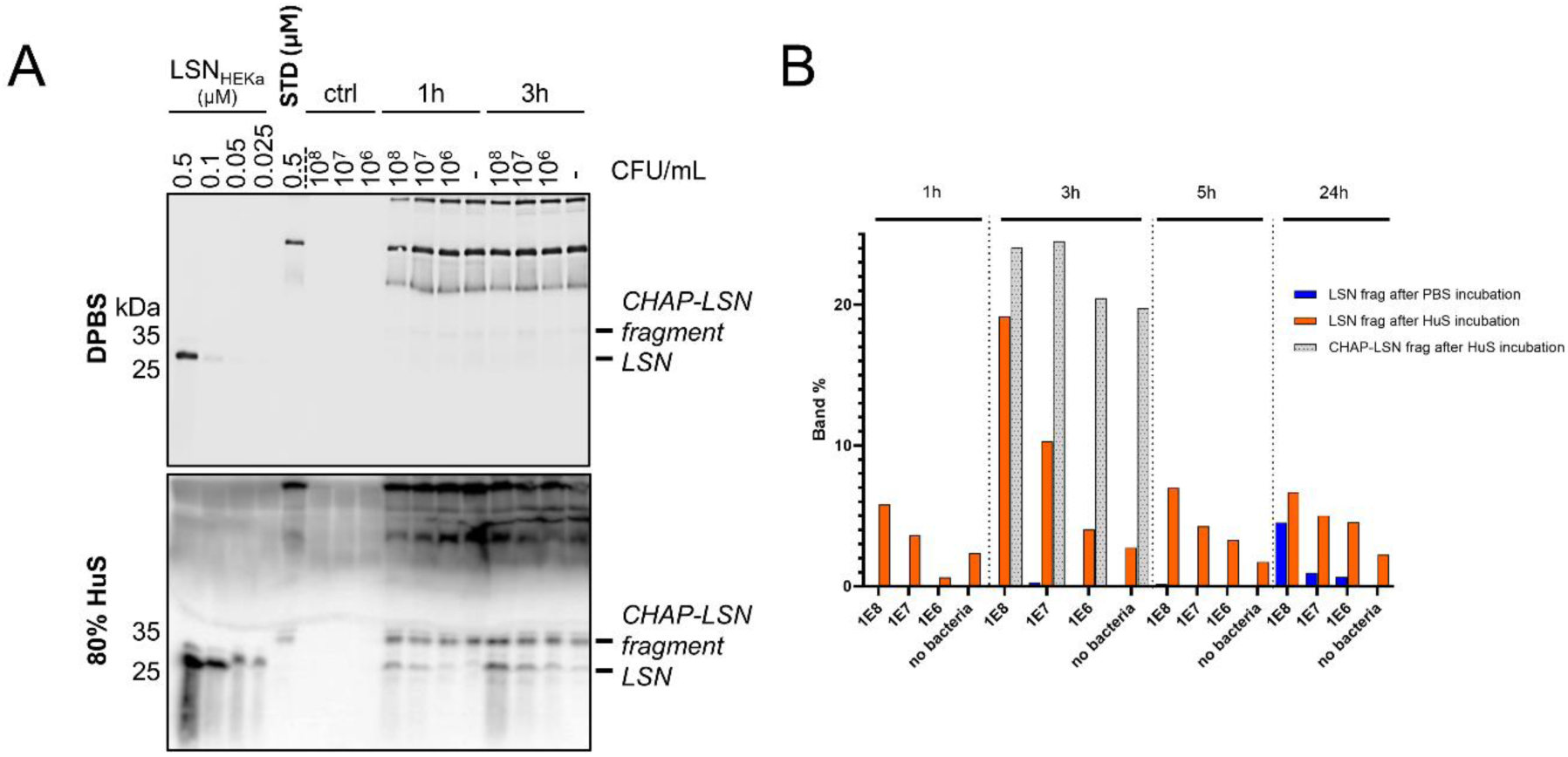
Dependency of LSN- and CHAP-LSN release on target cells and human serum. (A) Purified Fc-L1-CHAP_opt_-LSN_HEK_ (0.5 µM) was incubated in presence of DPBS (top) or 80% human serum (bottom) with increasing CFU/mL of *S. aureus* BAA-1717 at 37°C. Samples of each condition were taken after 3, 5 or 24 hours and were analyzed by SDS-PAGE, followed by Western blotting using a LSN-directed antibody for detection. (B) Quantification of full-length, CHAP-LSN fragment and LSN fragment in reactions shown in panel (A) in presence of DPBS (blue) and human serum (orange) after different incubation times. Protein amounts are expressed as percentages of the total anti-LSN signal in each lane of the western blot. Data for CHAP-LSN fragment (dotted gray) is only shown for the 3 h timepoint. Frag: fragment. STD: standard Fc-L1-CHAP_opt_-LSN_HEK_ at 0.5 µM.

Release of LSN in serum was accelerated by both intact cells and peptidoglycan and was dependent on a catalytically active PepM23 domain, since a catalytically inactive LSN variant (Fc-L1-CHAP-LSN*_HEK_) did not form any fragments (Figure 6A, B). Inactivation of the CHAP domain in an Fc-L1-CHAP*-LSN_HEK_ derivative prevented the formation of the CHAP-LSN fragment, but not the LSN fragment. In TKAs, Fc-L1-CHAP-LSN*_HEK_ was completely inactive whereas Fc-L1-CHAP*-LSN_HEK_ showed similar log10 CFU reductions compared to the parental Fc-CHAP-LSN construct (Figure 6C).

**Figure 6:**
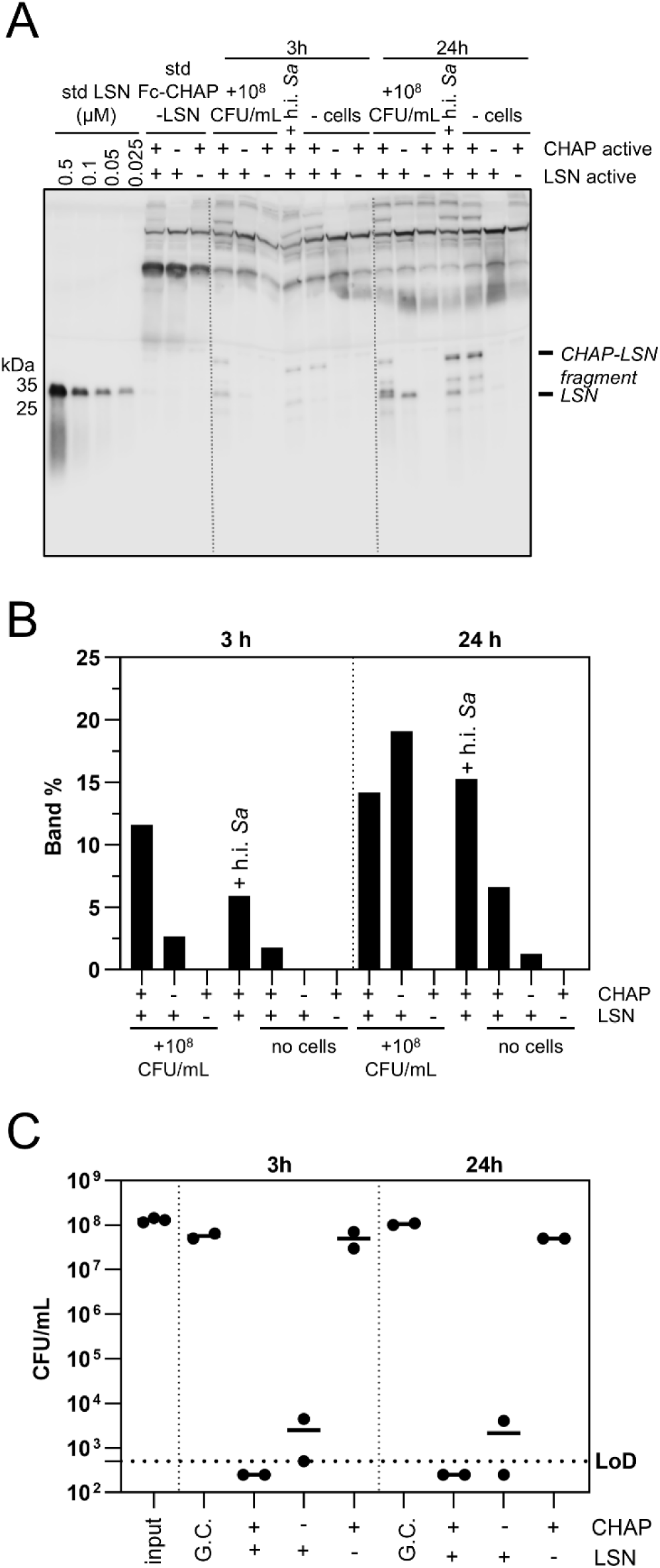
Dependency of LSN- and CHAP-LSN release on CHAP and PepM23 catalytic activity. (A) Purified Fc-L1-CHAP-LSN_HEK_ and variants thereof containing inactivating mutations in the CHAP domain (Fc-L1-CHAP*-LSN_HEK_) or PepM23 domain (Fc-L1-CHAP-LSN*_HEK_) were incubated in 80% human serum, in presence or absence of 10^8^ CFU/mL *S. aureus* BAA-1717 or in presence of a solution of heat inactivated *S. aureus* BAA-1717 (h.i. *Sa*, at an OD_600nm_ of 0.5) for 1, 3 and 24 h. Samples were analyzed for protein content by SDS-PAGE followed by western blotting. LSN_HEK_ was used as LSN standard in this experiment (B) Quantification of the released LSN bands shown in panel (A). Band intensities were integrated and expressed as a percentage intensity relative to the 0.5 µM recombinant lysostaphin standard loaded on each gel. (C) As in panel (A), but samples at the indicated timepoints were analyzed by quantitative plating for log10 CFU reduction versus a control lacking lysins. LOD: limit of detection. +: active domain, -: inactive domain

Taken together, these data suggested that LSN release occurs preferentially on the surface of bacteria, is catalyzed by the PepM23 domain of LSN and requires an unknown additional cofactor present in human serum, e.g. potentially Zn^2+^ or a serum protease. The data also suggested that the release of LSN contributes to the potency of Fc-CHAP-LSN constructs (as seen with Fc-L1-CHAP*-LSN_HEK_, which only generates an LSN fragment but is similarly active to the parental lysin), whereas generation of a CHAP-LSN fragment may or may not contribute to activity.

We hypothesized that target-cell dependent LSN-release from Fc-CHAP-LSN is due to the local enrichment of lysin on the peptidoglycan surface, which may increase the rates of encounter between PepM23 domains on one Fc-CHAP-LSN molecule and the respective cleavage sequence on the other Fc-CHAP-LSN molecule. To test this idea, we next asked if LSN-release could also be recapitulated by artificially localizing Fc-CHAP-LSN on the two-dimensional surface of an Expi293 cell, a scenario similar to lysin binding to the peptidoglycan surface of *S. aureus* which results in a drastic local increase in lysin concentration. For this purpose, we fused a flexible linker sequence followed by a PDGFR transmembrane domain to the C-terminus of Fc-CHAP-LSN, such that expression of this construct would lead to its anchoring to the plasma membrane facing towards the extracellular space (Figure 7A). Constructs encoding catalytically active and inactive Fc-CHAP-LSN variants Fc-L1-CHAP-LSN_HEK_-PDGFR, Fc-L1-CHAP_opt_-LSN_HEK_-PDGFR and Fc-L2-CHAP_opt_-LSN_HEK_-PDGFR were transfected into Expi293 cells, followed by expression and analysis of protein content in cell membranes by SDS-PAGE and western blotting (Figure 7B). Expression of the membrane-bound active Fc-L1-CHAP-LSN_HEK_-PDGFR, Fc-L1-CHAP_opt_-LSN_HEK_-PDGFR and Fc-L2-CHAP_opt_-LSN_HEK_-PDGFR, but not their catalytically inactive counterparts, resulted in the formation of a prominent fragment band corresponding to the LSN-PDGFR fusion (∼34 kDa). Direct comparison of LSN-fragmentation in secreted versus displayed lysins at different timepoints after transfection further demonstrated an increase in free LSN levels after lysin display (up to ∼12 fold versus secreted), confirming our notion that LSN-release is triggered by local lysin enrichment (Figure 7C). Interestingly, while the rates of LSN-release of displayed lysins were construct-independent, LSN-release rates in secreted constructs were strongly increased for constructs with reduced thermal stability and yield (i.e. for Fc-L1-CHAP-LSN_HEK_ and Fc-L1-CHAP_opt_-LSN_HEK_, Supplementary Table 1). This suggests that, for those constructs, the LSN cleavage site may be more accessible or the lysin may form more readily higher order aggregates in solution, promoting increased LSN-release even in the absence of target cells.

**Figure 7:**
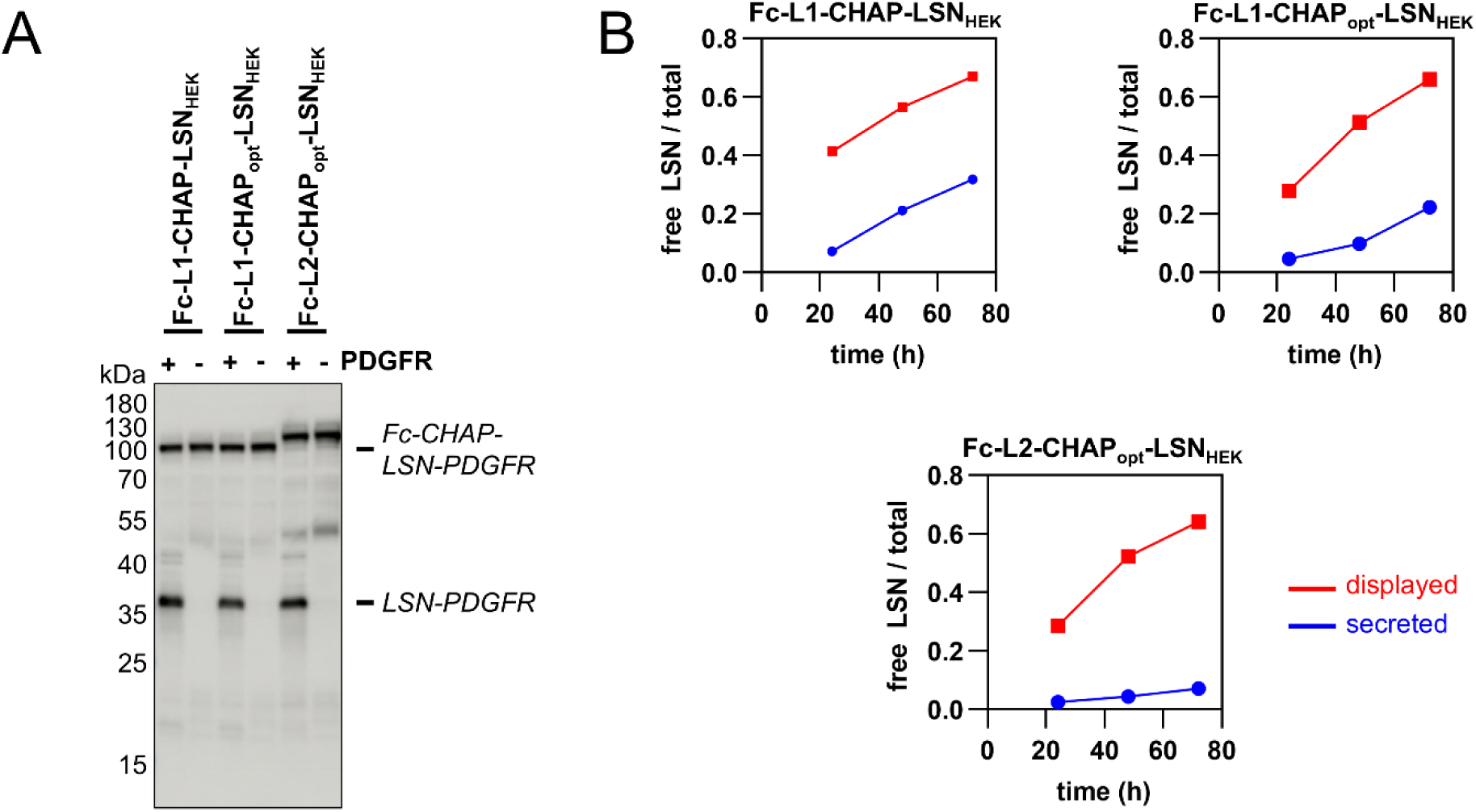
Proximity-induced LSN-release recapitulated on Expi293 plasma membranes. (A) PDGFR-fusions Fc-CHAP-LSN variants Fc-L1-CHAP-LSN_HEK_-PDGFR, Fc-L1-CHAP_opt_-LSN_HEK_-PDGFR, and Fc-L2-CHAP_opt_-LSN_HEK_-PDGFR containing catalytically active or inactive domains (Fc-L1-CHAP*-LSN*_HEK_-PDGFR, Fc-L1-CHAP*_opt_-LSN*_HEK_-PDGFR and Fc-L2-CHAP*_opt_-LSN*_HEK_-PDGFR) were transfected into EXPI293 cells. Cells were harvested and membrane bound protein species analyzed by SDS-PAGE followed by Western Blotting using an LSN-directed antibody for detection. Protein bands corresponding to full-length Fc-CHAP-LSN-PDGFR and the cleaved LSN-PDGFR fragment are indicated. (C) Time course of LSN-release in secreted versus displayed Fc-CHAP-LSN constructs. Fc-L1-CHAP-LSN_HEK_, Fc-L1-CHAP_opt_-LSN_HEK_ and Fc-L2-CHAP_opt_-LSN_HEK_ without C-terminally (blue, “secreted”) fused PDGFR domains and with fused PDGFR domain (red, “displayed”) were transfected into EXPI293 cells. Cells were harvested at the indicated times and supernatants or cellular fractions analyzed by SDS-PAGE followed by Western blotting using an LSN-directed antibody for detection of protein species. Bands corresponding to released LSN(-PDGFR) were integrated and divided by the total Western signal corresponding to full-length Fc-CHAP-LSN(-PDGFR), fragmented CHAP-LSN(-PDGFR) and LSN(-PDGFR).

**Figure 8:** SAXS analysis reveals an extended solution architecture of Fc-L1-CHAP_opt_-LSN_HEK_. (A) Solution model of the Fc-L1-CHAP_opt_-LSN_HEK_ dimer derived from SAXS rigid-body modeling. One trans-dimer configuration is shown as an illustrative example. The Fc region is shown in blue, CHAP in grey, PepM23 in light green, SH3 domains in light green, and the helical linker in dark green. The two CHAP-LSN arms adopt an extended arrangement, resulting in an inter-arm PepM23 - CHAP distance of ∼20 nm. (B) SAXS scattering data for Fc-L1-CHAP_opt_-LSN_HEK_ overlaid with the theoretical scattering curve calculated from the rigid-body model shown in (A). Experimental data and model fits show good agreement across the measured q-range (intensity in arbitrary units, A.U.). (B) (C) Surface representation of the PepM23 domain in PDB: 4LXC showing the distance between the N-terminus (Nt, red) and the catalytic pocket (formed by amino acids H32, D36, H115, side chains shown as red sticks) of the LSN PepM23 domain. The catalytic Zn^2+^ ion is shown as a grey sphere.

Based on these results, we hypothesized that LSN release is driven by self-cleavage within the CHAP–LSN linker, as the sequence KKA**GG**AAT resembles known LSN substrates and the *S. aureus* cross-bridge KAGGGGG (Antenucci et al., 2024). In agreement with this hypothesis, a Fc-L2-CHAP_opt_-LSN_HEK_ variant carrying a K143A mutation in the CHAP C-terminus (KAA**GG**AAT; Fc-L2a-CHAP_opt_var_-LSN_HEK_) exhibited enhanced cleavage in solution (Supplementary Fig. 5A and 5B). To map the truncation underlying the ∼50 kDa Fc-CHAP-LSN species, the corresponding Fc-L2a-CHAP_opt_var_-LSN_HEK_ fragment (Supplementary Fig. 5B) was analyzed by tryptic LC–MS/MS and corroborated with intact mass measurements. Toward the C terminus, peptide intensities dropped sharply, indicating loss of the distal C-terminal region in the truncated form (Supplementary Fig. 5C). Intact mass analysis revealed a degradation product at 50,933.7 Da post-deglycosylation, most consistent with a fragment spanning residues 1–467 bearing a sulfonic group (+48 Da). This aligns with extensive cysteine tri-oxidation observed for this protein, shown as main signal (Supplementary Fig. 5D and 5E). This assignment was further supported by the intact mass of the non-truncated species prior to deglycosylation, which closely matches the expected mass when a +48 Da modification and the observed glycan composition are considered (Supplementary Fig. 5D). Collectively, the mass spectrometry and mutagenesis data support a cleavage site in the KAAG/GAAT motif in the CHAP–LSN junction.

SAXS data collected for Fc-L1-CHAP_opt_-LSN_HEK_ were acquired and analyzed using rigid-body modeling to assess whether its solution architecture permits inter-monomer proteolytic cleavage within the dimer. Multiple independent CORAL reconstructions assuming P2 symmetry yielded dimeric models with elongated overall shapes and good agreement with the experimental scattering data (Figure 8A and B).

Two alternative dimer arrangements, referred to as trans- and cis-dimer conformations, were consistently obtained from the SAXS data and exhibited comparable fit quality. As the mechanistic conclusions drawn from the SAXS analysis are independent of the relative orientation of the two monomers, one trans-dimer model is used here solely as an illustrative example to discuss the spatial relationship between domains.

In both conformations, the Fc region mediates dimerization, while the CHAP-LSN arms extend outward from the Fc core, resulting in an overall elongated particle (Figure 8A). Measurement of inter-domain distances shows that the catalytic CHAP and PepM23 domains of the two CHAP-LSN arms are separated by approximately ∼20 nm via the rigid linkers, a structure that is incompatible with inter-monomer proteolytic cleavage within the dimer.

Consistent with this, inspection of the LSN crystal structure (PDB: 4LXC) (Sabala et al., 2014) indicates that the N-terminus of LSN is physically occluded and positioned approximately ∼2.5 nm distant from the PepM23 active site within a single monomer, rendering intra-monomer cleavage geometrically unfavorable (Figure 8C). Thus, the SAXS-derived models support an extended Fc-CHAP-LSN architecture in which neither intra- nor inter-monomer CHAP-mediated cleavage is structurally supported.

Taken together, these data indicate that inter-molecular LSN self-release occurs when LSN is anchored to the cell-surface, presumably driven by the high local enzyme and substrate concentrations.

## Discussion

This is the first report to show that half-life extension of LSN via human IgG1 Fc is feasible. However, this architecture requires a second catalytic domain (CHAP) to retain LSN’s broad staphylococcal strain coverage. The resulting Fc-CHAP-LSN is highly active against both planktonic cells and biofilms, and *in vivo,* in *S. aureus* sepsis models, with the activity being mainly driven by the LSN catalytic domain.

Additionally, we provide compelling evidence for an elegant mechanism of action for Fc-CHAP-LSN: it acts both as an enzyme and a substrate to release free LSN (Figure 9), the process being enhanced by locally concentrating enzyme and the substrate, e.g. via target engagement (Figure 9A) or surface co-localization (Figure 9B). The release of LSN correlates with bactericidal activity, and both release and bactericidal activity require a catalytically active PepM23 domain. The LSN cleavage site has been mapped to the Gly-Gly region in the CHAP-LSN linker by mass spectrometry; neighboring residues to the Gly-Gly site play a role in modulating LSN release activity and it is envisaged that sequences with better targeted release can be obtained by linker mutagenesis.

**Figure 9:** Proposed mechanism of LSN-release from Fc-CHAP-LSN. (A) LSN-release triggered by target cell binding. Binding of full-length Fc-CHAP-LSN to the peptidoglycan layer of target *S. aureus* cells locally enriches the protein, resulting in strongly increased release of the LSN moiety from one lysin molecule by the LSN-domain from a second lysin molecule. Due to its small size, released LSN can efficiently penetrate and hydrolyze the peptidoglycan layer. (B) LSN-release recapitulated on EXPI293 membranes. Similar to (A), the high local concentration of displayed lysin promotes trans-lysin cleavage of LSN, resulting in membrane bound LSN-PDGFR fragments.

LSN release does not occur (or occurs at a much slower rate) in solutions of Fc-CHAP-LSN because both enzyme and substrate are relatively diluted. This agrees with previous studies which report a Km in the mM range for LSN with synthetic poly-glycine peptides (Grishin et al., 2020). It also does not occur intramolecularly, presumably because: 1. the access of the catalytic site of the PepM23 domain to the short Gly-Gly CHAP-PepM23 linker within the same monomer is sterically hindered in a fully folded PepM23 domain and 2. the inter-arm cleavage (of CHAP-PepM23 linker from one monomer by the PepM23 catalytic site from the second monomer in the Fc-dimer) is prevented by the rigid nature of the helical Fc-CHAP linker. Hence, optimization of the targeted (vs non-targeted) LSN release kinetics can be achieved by engineering both the CHAP-LSN as well as the Fc-CHAP linker sequences.

Larger fragments are generated during Fc-CHAP-LSN expression in mammalian cell supernatants, corresponding to cleavage in poly-Gly sequences in the CHAP domain, but their intensity decreases with increasing lysin stability, indicating that partial unfolding occurs in the CHAP domain in this architecture. This is supported by thermal stability measurements: Tm1 for Fc-L1-CHAP-LSN_HEK_ is 45.1 C (interestingly for the CHAP inactive variant Fc-L1-CHAP*-LSN_HEK_ Tm1 is 57.7 C, Supplementary Table 1), while for the corresponding LysM-CHAP variant (L2-1 in Badarau et al., 2026) Tm1 is 53.9 C. There is also some basal LSN release from Fc-CHAP-LSN, observed in mammalian cell supernatants and in presence of human serum, i.e. in absence of target cells, which appears to correlate with protein stability (Figures 5 - 7), indicating that misfolding/aggregation can also trigger LSN release.

The mechanism presented here can be exploited for the half-life extension of other peptidase-based bacterial lysins, counteracting the negative impact on activity brought about by the increased size and avidity of the Fc fusion protein (Badarau et al., 2026). It can also be exploited for other peptidase-based therapeutics, targeting surface presented or aggregated substrates, or for local release of peptide-linked payloads.

## Conflict of Interest

ZV, DK, BB, RQ, SDM, AMH, JS, RB, KK, MM, MZ, MF and AB are employees of BioNTech and hold shares in the company. LC is a former employee of BioNTech and holds shares in the company. ZV, DK, BB, RQ, SDM, MM, AB and LC are inventors on patent applications related to the technology described in the manuscript.

## Author Contributions

AB conceptualized and directed the research. ZV directed the in vitro functional characterization and in vivo work and the mechanistic work in presence of bacterial cells. DK and BB led the protein engineering work. RQ led the protein production and biophysical characterization work. SDM and JS performed and analyzed the spike-in experiments. AMA designed and coordinated the *in vivo* studies. RB performed and analyzed the lysin degradation experiments in presence of bacterial cells and the lytic assays. KK, MM and MF performed and analyzed the Expi293 display experiments. MZ performed and analyzed the bacterial growth inhibition experiments. MF produced and characterized the recombinant proteins. LC contributed to conceptualization and direction of the research, reviewed data and progress. All authors contributed to data interpretation. ZV, DK, BB and AB wrote the manuscript with input from RQ. All authors reviewed the manuscript.

## Acknowledgments

We would like to thank Philipp Czermak for support with protein production and to Gurli Wertanek, Manuel Maluenda and Sruthi Kurugodu for their excellent technical work. We thank Fiona Powell and Amanda Gallagher for critical review of the manuscript. We acknowledge Evotec UK (Ltd) for performing the animal studies, the Vienna BioCenter core facilities (Protein Technologies) for protein production, The Mass Spectrometry Facility of the MFPL for the mass spectrometry data acquisition and analysis, and BioSAXS for the recording and analysis of SEC-SAXS data.

## Data Availability Statement

SAXS raw data has been submitted for deposition in the SASBDB (accession code pending)

