## Supplemental Material for "A chimeric, half-life extended lysin with a unique mode of action"

### *Supplementary Material*

#### Supplementary method

*Small-scale expression and activity testing in supernatant*

*Transfection and harvest.*

DNA concentrations were normalized to 100 ng/µL. DNA was mixed with transfection reagents, followed by transfer onto 1 mL Expi293 cultures in 24 well format. Transfections, enhancer addition and cell culture conditions were as in the main text. Supernatants were finally harvested (300 × g, 5 minutes), aliquoted and supplementation with 0.5 0M Tris(2-carboxyethyl)phosphine (TCEP). Samples were either processed directly or frozen at -80 °C.

*High throughput growth inhibition assay in supernatants.*

Overnight culture of *Staphylococcus. aureus* ATCC 43300 was grown in tryptic soy broth (TSB) at 37 °C and shaking at 220 rpm, diluted 1:20 into TSB to an optical density at 600 nm ($OD_{600}$) of ~0.05, and further grown until reaching mid-log phase ($OD_{600}$ ~1.0). This culture was then diluted in cation-adjusted Mueller–Hinton broth (caMHB) supplemented with 20% heat-inactivated horse serum to reach a final inoculum of ~5 × $10^5$ CFU/mL. A 2-fold dilution series of the supernatant was prepared in caMHB supplemented 20% heat-inactivated horse serum. Reactions were set up in 96-well U-bottom plates by mixing 30 µL of supernatant solution with 70 µL bacterial suspension. Plates were statically incubated at 37 °C for 17-19 h. Minimal inhibitory concentration (MIC) was defined as the lowest concentration of lysin that still prevented growth by visual inspection.

*OD reduction assays.*

*S. aureus* BAA-1717 was cultured aerobically on Columbia Agar with 5% sheep blood or in TSB at 37 °C. Pre-cultures were prepared by inoculating TSB with a few bacterial colonies grown on an agar plate, followed by overnight incubation. The pre-culture was used to inoculate 100 mL of TSB at an initial $OD_{600}$ of 0.05. Cells were grown in baffled flasks at 37°C and 130 rpm until an $OD_{600}$ of ~1.1 was reached. The bacterial cells were subsequently harvested by centrifugation (4000 × g, 10 min). Cells were washed twice in DPBS supplemented 1 mM $CaCl_2$ and 0.5 mM $MgCl_2$, followed by resuspension in the same buffer at a final $OD_{600}$ of 2. Supernatants were diluted 1:8 to 1:16 in Expi293 expression medium supplemented with 0.5 mM TCEP. 100 µL of the diluted supernatants were transferred into a flat-bottomed 96-well plate and mixed with an equal volume of the bacterial suspension, yielding a final $OD_{600}$ of 1. The reaction plate was immediately placed into a Spectramax Plus plate reader (Molecular Devices) pre-heated to 37 °C. $OD_{600}$ was measured at intervals of 20 s over a period of 2 h. Rates of OD reduction were extracted as the maximum slope of the OD reduction curves. To mitigate the impact of noise in the OD data, which can hinder accurate determination of the maximum slope, the data was smoothed using a cubic spline representation. The maximum slope was identified as the minimum of the first derivative of the spline. Data smoothing and slope determination were implemented in Python using the SciPy package and the scipy.interpolate.make_smoothing_spline function with the default lambda parameter.

#### Supplementary Figures and Tables

#### Supplementary Figures

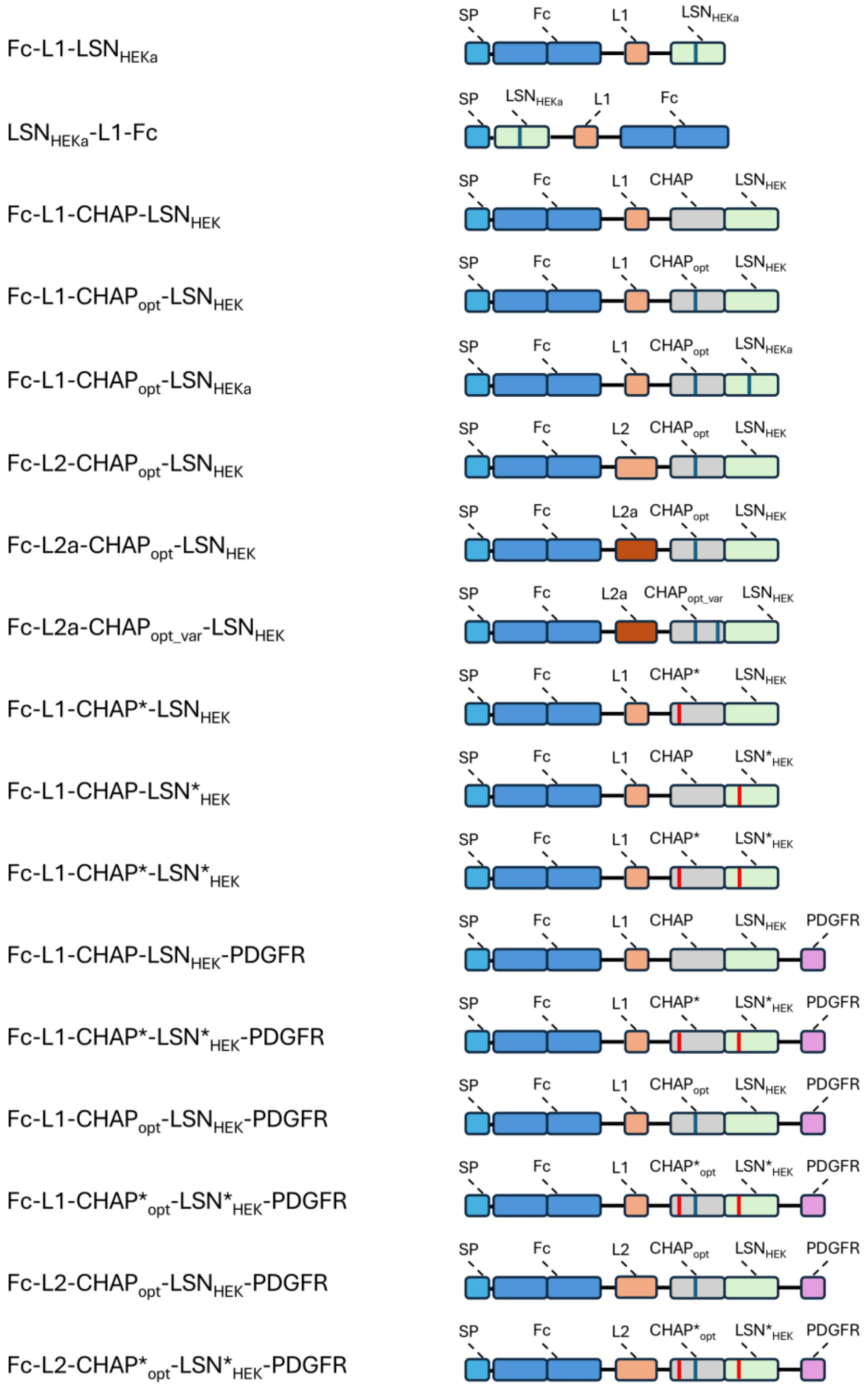


**Supplementary Figure 1. Graphical representation of the diFc-CHAP-LSN constructs (only one monomer shown).** Each construct consists of a signal peptide (SP) combined with purification tags, Fc domain (FC), a CHAP domain and LSN domain. The SP represents the murine Ig κ light-chain signal peptide, a TG dipeptide, an N-terminal hexahistidine tag, and a

HRV 3C protease cleavage site. The Fc represents the IgG1 Fc domain. L1 refers to the helical linker with 8 repeats while L2 and L2a refer to the helical linkers with 14 repeats plus a 12 amino acid serum stable peptide. L2 and L2a only differ in the composition of the 12 amino acid peptide. CHAP refers to the CHAP domain of L14DR, $CHAP_{opt}$ is the CHAP domain of L14DR with an additional Y46D mutation and $CHAP_{opt\_var}$ contains an additional K143A mutation. Inactive variants are marked with an asterisk (CHAP: C27S; LSN: H82A or H113A). Display constructs contain a C-terminal PDGFR transmembrane domain.

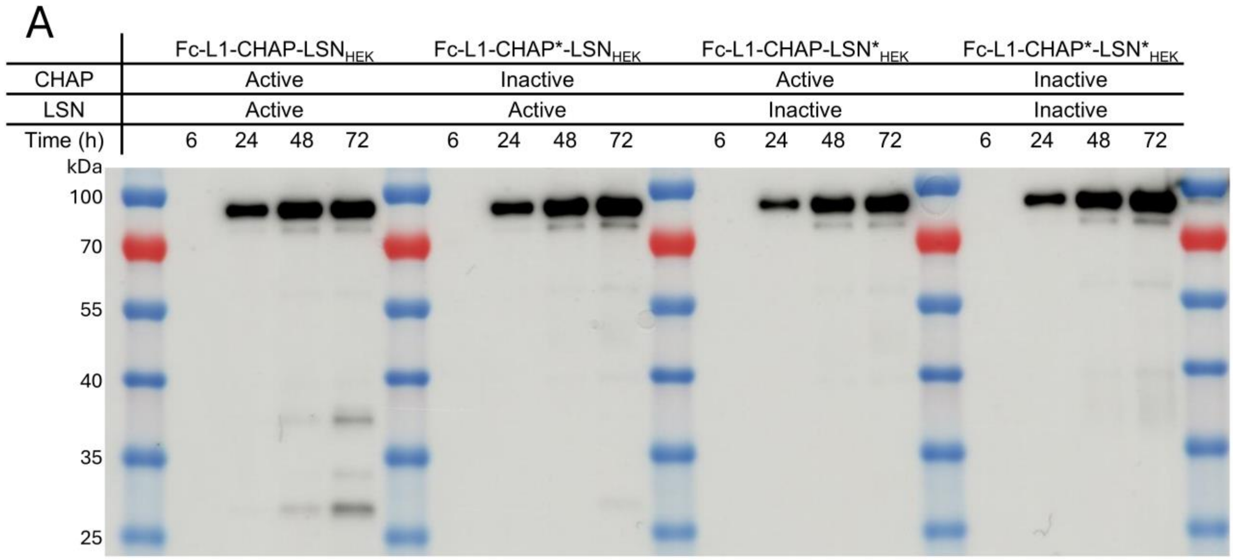


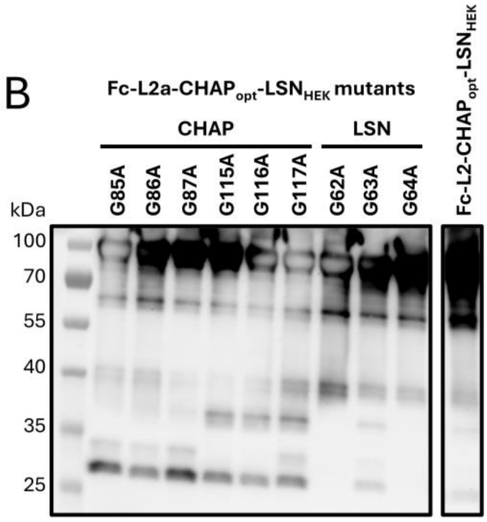


C

| Construct | Secretion (µg/ml) | ORA rate (OD/h) | MIC (µg/mL) |
|---|---|---|---|
| Fc-L2-CHAP$_{opt}$-LSN$_{HEK}$ | 126 | -0.506 | 1.18 |
| Fc-L2a-CHAP$_{opt\text{-}G85A}$-LSN$_{HEK}$ | 101 | -0.749 | 1.89 |
| Fc-L2a-CHAP$_{opt\text{-}G86A}$-LSN$_{HEK}$ | 90 | -0.617 | 0.84 |
| Fc-L2a-CHAP$_{opt\text{-}G87A}$-LSN$_{HEK}$ | 62 | -0.909 | 1.16 |
| Fc-L2a-CHAP$_{opt\text{-}G115A}$-LSN$_{HEK}$ | 88 | -0.525 | ≤0.41 |
| Fc-L2a-CHAP$_{opt\text{-}G116A}$-LSN$_{HEK}$ | 98 | -0.469 | 0.92 |
| Fc-L2a-CHAP$_{opt\text{-}G117A}$-LSN$_{HEK}$ | 81 | -0.663 | ≤0.38 |
| Fc-L2a-CHAP$_{opt}$-LSN$_{HEK\text{-}G62A}$ | 143 | -0.085 | 42.88 |
| Fc-L2a-CHAP$_{opt}$-LSN$_{HEK\text{-}G63A}$ | 99 | -0.333 | 14.85 |
| Fc-L2a-CHAP$_{opt}$-LSN$_{HEK\text{-}G64A}$ | 110 | -0.222 | n.d. |

**Supplementary Figure 2. Western blot analysis of diFc-CHAP-LSN variants in supernatant.** (A) Western blot of a time-course secretion experiment in which Fc-L1-CHAP-LSN$_{HEK}$ is compared to the Fc-L1-CHAP*-LSN$_{HEK}$, Fc-L1-CHAP-LSN*$_{HEK}$ and Fc-L1-CHAP*-LSN*$_{HEK}$. Aliquots were taken after 6, 24, 48 and 72 hours. Where the active variant results in the appearance of three fragmentation bands which correspond to the release of LSN (27 kDa) and two CHAP-LSN fragments (33 and 37 kDa). A faint LSN band can be observed in the Fc-L1-CHAP*-LSN$_{HEK}$ sample around 27 kDa. For all constructs faint fragmentations bands can be observed at 40 and 57 kDa, assumed to be independent of the catalytic activity of the constructs. (B) Western blot of the supernatant samples in which single glycine were substituted within the GGG stretches of Fc-L2a-CHAP$_{opt}$-LSN$_{HEK}$ to alanine. Six CHAP variants (CHAP$_{opt\text{-}G85A}$, CHAP$_{opt\text{-}G86A}$, CHAP$_{opt\text{-}G87A}$, CHAP$_{opt\text{-}G115A}$, CHAP$_{opt\text{-}G116A}$ or CHAP$_{opt\text{-}G117A}$) and three LSN variants (LSN$_{HEK\text{-}G62A}$, LSN$_{HEK\text{-}G63A}$ or LSN$_{HEK\text{-}G64A}$) were generated. The western blot was overloaded to reveal the presence of truncation products. Four lower molecular truncation products can be observed (27 – 33 – 37 – 40 kDa). Point mutations in the CHAP$_{opt\text{-}85GGG87}$ stretch result in the loss of signal of the 37 kDa band, while the point mutations in the CHAP$_{opt\text{-}115GGG117}$ stretch result in the loss of signal of the 33 kDa band. The mutations in LSN result in severely reducing the activity of the constructs, less to no degradation bands are observed at 27 kDa, 33 kDa or 37 kDa. The closest related reference sample (Fc-L2-$CHAP_{opt}$-$LSN_{HEK}$) on the same gel is shown as a reference. (C) Secretion yield and activity data of the mentioned constructs in supernatant. Secretion yields were quantified by SDS-PAGE, OD reduction assay rate (ORA rate) was performed with *S. aureus* BAA-1717 and the MIC was determined for *S. aureus* ATCC 43300.

.

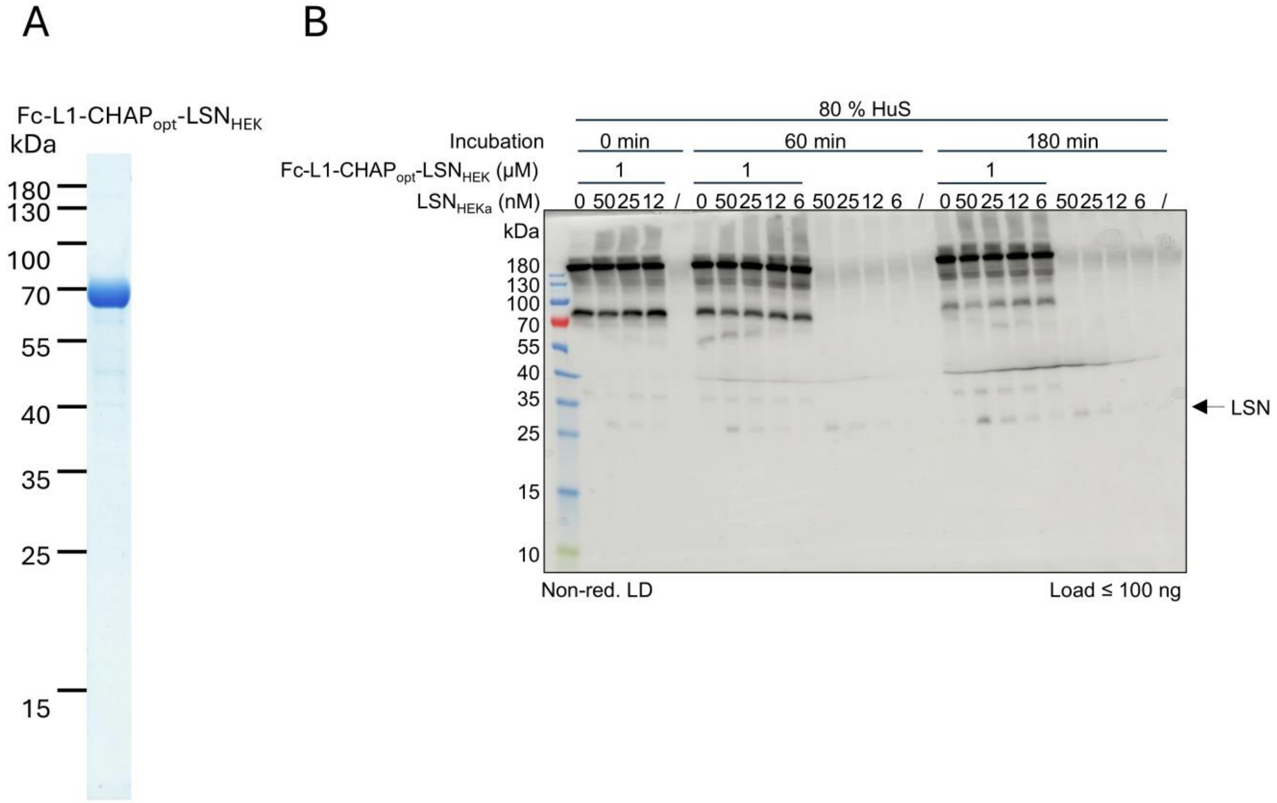


**Supplemental Figure 3:** (A) SDS–PAGE analysis of purified Fc-L1-CHAP$_{opt}$-LSN$_{HEK}$ under reducing conditions. The protein sample (2.5 µg) were resolved on an AnykD Criterion™ TGX precast gel and visualized by InstantBlue® protein stain. Molecular weight markers (kDa) are indicated. A predominant band at the expected molecular weight is observed, consistent with high sample purity. (B) LSN-induced LSN-release from Fc-L1-CHAP$_{opt}$-LSN$_{HEK}$. Western blot corresponding with the data from Figure (4). Purified LSN$_{HEKa}$ was incubated at increasing concentrations in 80% human serum and in presence of $10^8$ CFU/mL *S. aureus* ATCC 43300 with or without a fixed concentration of Fc-L1-CHAP$_{opt}$-LSN$_{HEK}$ (1 µM) for up to 3h at 37°C. Samples were analyzed by SDS-PAGE followed by Western Blotting to quantify the amounts of free LSN in each reaction.

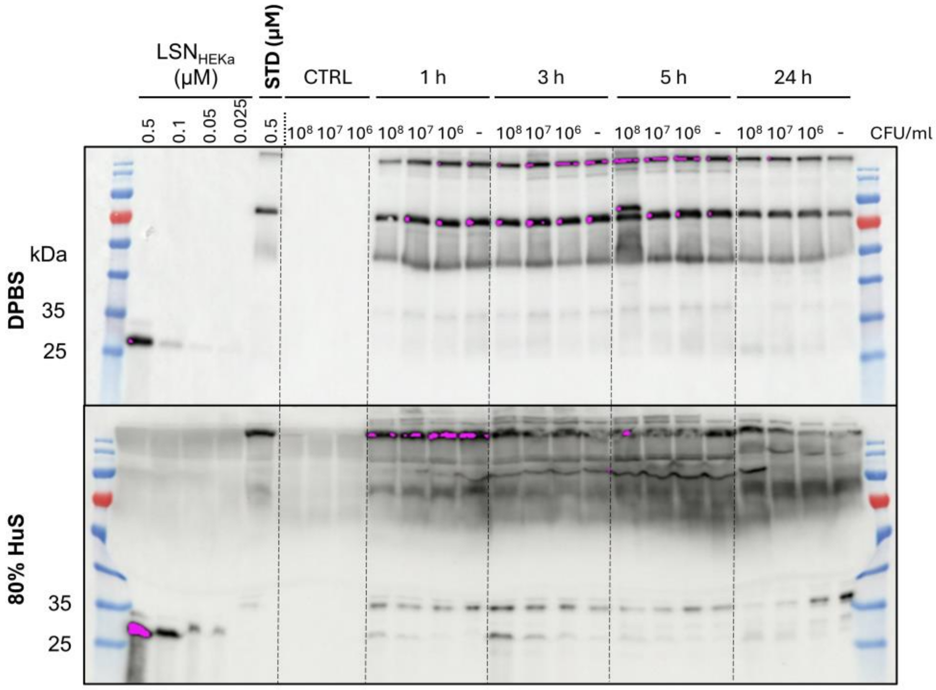


**Supplementary Figure 4.** Full western blot used for quantification in Figure 5. Purified Fc-L1-CHAP$_{opt}$-LSN$_{HEK}$ (0.5 µM) was incubated in presence of DPBS (top) or 80% human serum (bottom) with increasing CFU/mL of *S. aureus* BAA-1717 at 37°C. Samples of each condition were taken after 3, 5 or 24 hours and were analyzed by SDS-PAGE, followed by Western blotting using a LSN-directed antibody for detection. If no bacteria were added the lane was denoted with "-". STD: standard Fc-L1-CHAP$_{opt}$-LSN$_{HEK}$ at 0.5 µM. CTRL: Control sample with bacteria only at different CFU/mL.

A

```
1   GPEPKSSDKT HTCPPCPAPE LLGGPSVFLF PPKPKDTLMI
41  SRTPEVTCVV VDVSHEDPEV KFNWYVDGVE VHNAKTKPRE
81  EQYNSTYRVV SVLTVLHQDW LNGKEYKCKV SNKALPAPIE
121 KTISKAKGQP REPQVYTLPP SRDELTKNQV SLTCLVKGFY
161 PSDIAVEWES NGQPENNYKT TPPVLDSDGS FFLYSKLTVD
201 KSRWQQGNVF SCSVMHEALH NHYTQKSLSL SPGKAEAAAK
241 EAAAKEAAAK EAAAKEAAAK EAAAKEAAAK ALEAEAAAKE
281 AAAKEAAAKE AAAKEAAAKE AAAKEAAAKA KPETVSGNEK
321 SNSGSSTLNY LKSLEGKGWD FDGSFGWQCF DLVNVYWNHL
361 YGHGLKGDYA KDIPYANNFN NEAKIYKNTP TFKAEPGDLV
401 VFSGRYGGGY GHTAIVLNGN YDGKLQKFQS LDQNWYGGGE
441 RKAEVAHKVV HNYENDMIFI RPFKAAGGAA THEHSAQWLN
481 NYKKGYGYGP YPLGINGGMH YGVDFFMNIG TPVKAISSGK
521 IVEAGWSNYG GGNQIGLIEN DGVHRQWYMH LSKYNVKVGD
561 YVKAGQIIGW SGSTGYSTAP HLHFQRMVNS FSNPTAQDPM
601 PFLKSAGYGK AGGTVTPTPN TGWKTNKYGT LYKSESASFT
641 PNTDIITRTT GPFRSMPQSG VLKAGQTIHY DEVMKQDGHV
681 WVGYTGNSGQ RIYLPVRTWQ KSTNTLGVLW GTIK
```

B

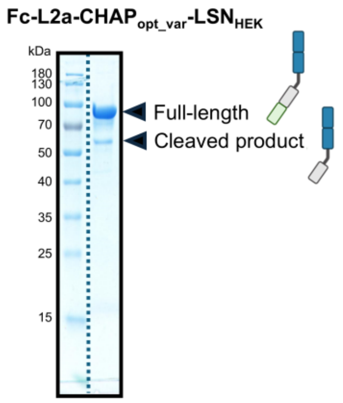


C

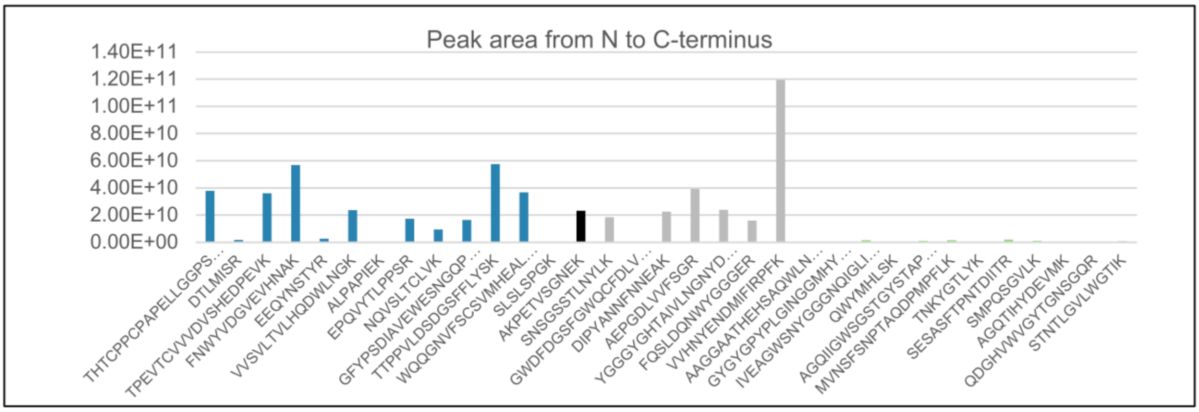


D

| Experimental masses | Delta mass (Da) | Interpretation | Mass error (Da) |
|---|---|---|---|
| 77952.0 | 58.6 | FL + deamidation + 57 Da | |
| 78140.8 | 247.4 | FL + deamidation + HexNac + 43 Da | |
| 78600.1 | 706.7 | FL + deamidation + Hex HexNAc NeuAc + 49 Da | |
| 78885.2 | 991.8 | FL + deamidation + Hex HexNAc NeuAc(2) + 43 Da | |
| 79068.5 | 1175.1 | etc | |
| 79556.5 | 1663.1 | | |
| 79826.7 | 1933.3 | | |

E

| Experimental masses | Delta mass (Da) | Interpretation | Mass error (Da) |
|---|---|---|---|
| 78213.2 | 205.6 | FL+ deamidation + HexNAc | + 1.4 |
| 78957.6 | 950.0 | FL + deamidation + Hex HexNAc NeuAc(2) | + 1.2 |
| 79208.6 | 1201.0 | ? | |
| 79905.4 | 1897.8 | FL + deamidation + 2(Hex HexNAc NeuAc(2)) | + 1.2 |

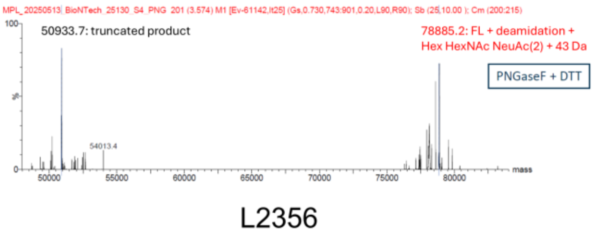


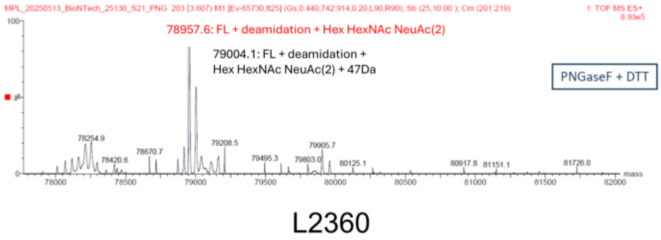


**Supplementary Figure 5. Mass spectrometry–based mapping of the cleavage site in variant Fc-L2a-CHAP$_{opt_var}$-LSN$_{HEK}$. (A)** Amino acid sequence of Fc-L2a-CHAP$_{opt_var}$-LSN$_{HEK}$ shown with residue numbering. The Fc region is shown in blue, the helical linker in black, CHAP in grey, PepM23 in light green, SH3 domains in light green. The proposed cleavage site GG is indicated in red. **(B)** Representative SDS-PAGE analysis of purified Fc-L2a-CHAP$_{opt_var}$-LSN$_{HEK}$ showing the full-length protein and a stable ~50 kDa cleavage product. Schematic cartoons illustrate the domain architecture of the full-length construct and the truncated species. **(C)** Peptide abundance profile along the Fc-L2a-CHAP$_{opt_var}$-LSN$_{HEK}$ sequence derived from FragPipe analysis of the 50 kDa gel slice. Following tryptic digestion and LC–MS/MS, peptide peak areas were extracted using FragPipe (24.0) and plotted from the N to the C-terminus. Only fully tryptic peptides without missed cleavages were included. Peptides C-terminal to Lys464 exhibit strongly reduced signal intensities relative to upstream peptides, despite the predominance of full-length protein in the sample, consistent with a C-terminal truncation occurring at the CHAP–LSN junction. **(D)** Intact mass analysis of PNGase F treated Fc-L2a-CHAP$_{opt_var}$-LSN$_{HEK}$ showing a dominant molecular species with a mass shift of approximately +43 Da relative to the theoretical full-length sequence mass and a distinct truncated species (50933.7 Da). **(E)** Intact mass analysis of PNGase F treated Fc-L2-CHAP$_{opt}$-LSN$_{HEK}$ showing the non-oxidized full-length form as the main peak, accompanied by satellite peaks with mass shifts of approximately +42 to +48 Da.

**Supplementary Table 1. Thermal stability and production yield of lysin constructs in three different formulation buffers.**

Thermal unfolding temperatures (Tm) and production yields are reported as mean ± SD across independent production batches for each lysin construct. Data are grouped by formulation buffer as follows: MES buffer (50 mM MES, pH 6.5, 300 mM NaCl, 1 mM $CaCl_2$, 5% glycerol, 50 mM arginine, 2 mM DTT), Histidine buffer (100 mM L-histidine, pH 6.5, 300 mM NaCl, 1 mM $CaCl_2$, 5% glycerol), and HEPES buffer (20 mM HEPES, 150 mM NaCl, pH 7.0).

| **MES buffer** | | | | | | |
|---|---|---|---|---|---|---|
| **Lysin** | **n (Tm)** | **Tm mean (°C)** | **Tm SD (°C)** | **n (Final yield)** | **Final yield mean (µg/mL)** | **Final yield SD (µg/mL)** |
| Fc-L1-CHAP-$LSN_{HEK}$ | 5 | 52.5 | 3.2 | 7 | 13.0 | 9.8 |
| Fc-L1-$CHAP_{opt}$-$LSN_{HEK}$ | 1 | 53.1 | NA | 3 | 66.0 | 35.8 |
| Fc-L2-$CHAP_{opt}$-$LSN_{HEK}$ | 1 | 54.6 | NA | 1 | 84.0 | NA |
| Fc-L2a-$CHAP_{opt}$-$LSN_{HEK}$ | 1 | 55.8 | NA | 1 | 34.4 | NA |
| **Histidine buffer** | | | | | | |
| **Lysin** | **n (Tm)** | **Tm mean (°C)** | **Tm SD (°C)** | **n (Final yield)** | **Final yield mean (µg/mL)** | **Final yield SD (µg/mL)** |
| Fc-L1-$CHAP_{opt}$-$LSN_{HEK}$ | 4 | 52.0 | 3.1 | 2 | 28.2 | 24.1 |
| Fc-L2-$CHAP_{opt}$-$LSN_{HEK}$ | 2 | 53.4 | 0.5 | 1 | 14.8 | NA |
| Fc-L2a-$CHAP_{opt}$-$LSN_{HEK}$ | 1 | 53.6 | NA | 0 | NA | NA |
| **HEPES buffer** | | | | | | |
| **Lysin** | **n (Tm)** | **Tm mean (°C)** | **Tm SD (°C)** | **n (Final yield)** | **Final yield mean (µg/mL)** | **Final yield SD (µg/mL)** |
| Fc-L1-CHAP-$LSN_{HEK}$ | 1 | 45.1 | NA | 6 | 9.5 | 6.8 |
| Fc-L1-CHAP*-$LSN_{HEK}$ | 1 | 57.7 | NA | 1 | 61.0 | NA |
| Fc-L1-$CHAP_{opt}$-$LSN_{HEKa}$ | 1 | 50.6 | NA | 4 | 48.4 | 16.4 |
| Fc-L1-CHAP-$LSN*_{HEK}$ | 1 | 55.1 | NA | 1 | 11.7 | NA |
